# Deconvolution Of Hematopoietic Stem/Progenitor Cell Signaling Predicts Inflammatory Niche Remodeling To Be A Determinant Of Tissue Failure And Outcome In Human AML

**DOI:** 10.1101/2023.05.05.539525

**Authors:** Lanpeng Chen, Eline Pronk, Claire van Dijk, Yujie Bian, Jacqueline Feyen, Tim van Tienhoven, Meltem Yildirim, Paola Pisterzi, Madelon de Jong, Alejandro Bastidas, Remco Hoogenboezem, Chiel Wevers, Eric Bindels, Bob Löwenberg, Tom Cupedo, Mathijs A. Sanders, Marc H.G.P. Raaijmakers

## Abstract

Cancer initiation is orchestrated by interplay between tumor-initiating cells and their stromal/immune environment. Here, by adapted scRNAsequencing, we decipher the predicted signaling between tissue-resident hematopoietic stem/progenitor cells (HSPCs) and their neoplastic counterparts with their native niches in the human bone marrow. LEPR^+^ stromal cells are identified as central regulators of hematopoiesis through predicted interactions with all cells in the marrow. Inflammatory niche remodeling and the resulting deprivation of critical HSPC regulatory factors is predicted to repress distinct high-output HSC subsets in *NPM1*-mutated AML, with relative resistance of clonal cells. Stromal gene signatures reflective of niche remodeling are associated with reduced relapse rates and favorable outcome after chemotherapy, across all genetic risk categories. Elucidation of the intercellular signaling defining human AML, thus, predicts that inflammatory remodeling of stem cell niches drives tissue repression and clonal selection, but may pose a vulnerability for relapse-initiating cells in the context of chemotherapeutic treatment.

**statement of significance:** Tumor-promoting inflammation is considered an enabling characteristic of tumorigenesis, but mechanisms remain incompletely understood. By deciphering the predicted signaling between tissue-resident stem cells and their neoplastic counterparts with their environment, we identify inflammatory remodeling of stromal niches as a determinant of normal tissue repression and clinical outcome in human AML.

**Key points:**

- A comprehensive taxonomy of the predicted interactions between LEPR^+^ stromal niches, HSPCs and adaptive/innate immune cells in the human NBM.
- Inflammation-associated decline of stromal niches in AML represses residual normal hematopoiesis with relative resistance of leukemic cells.
- Inflammatory decline of stromal niches is associated with reduced relapse risk and favorable outcome.

## INTRODUCTION

Cancer initiation and drug resistance are orchestrated by an interplay between tumor initiating cells, residual normal tissue stem and progenitor cells and ancillary tissue-resident cells, comprising the stromal and immune environment [1]. Ultimately, insight into the signaling cascades operative between all these cells in a neoplastic state is required to obtain a comprehensive appreciation of the complexity of tumorigenesis and tumor survival in the context of therapy.

One important component of all human tissues are stromal cells. Bone marrow stromal cells (BMSC) pervade the tissue in extensive networks that are estimated to comprise up to 20% of the marrow’s cellular volume and associate with the vast majority of hematopoietic cells [2]. They represent a likely heterogeneous population of cells, with a subset considered critical for the maintenance and homeostatic regulation of hematopoietic stem and progenitor cells (HSPCs) [3]. A subset of stromal cells comprising the HSPC niche has been characterized in mice by expression of the leptin receptor (LEPR) [4, 5] and high expression of CXCL12 (hence dubbed ‘CXCL12- abundant reticular (CAR) cells’ [6, 7] located in the vicinity of sinusoid vessels [2, 5, 8]. These stromal cells are major sources of stem cell factor (SCF) and interleukin-7 and considered critical regulators of HSCs, multipotent progenitors, lymphoid progenitors as well as natural killer, B and plasmacytoid dendritic cell development [4, 9–12].

It is, however, important to emphasize that these insights have been derived mostly from genetic studies in non-human species, in particular mice. The biology, heterogeneity and interactions of LEPR^+^ stromal cells in the human bone marrow have remained largely elusive, in part because it is difficult to retrieve these cells in sufficient numbers, likely because they reside along fibers of extracellular matrix, limiting their capture by aspiration.

Insight into the biology of human LEPR^+^ stromal cells and in particular their interaction with HSPCs, is of likely further relevance to the biology of myeloid neoplasms, including acute myeloid leukemia (AML). AML is caused by genetic events occurring in HSPCs, but mouse modeling and humanized *ex vivo* modeling have implicated stromal niche alterations in the initiation, maintenance, drug resistance and progression of AML [13–23]. Transcriptional alterations in *Lepr*+ stromal cells, including the downregulation of hematopoietic regulatory factors, have further been documented in mice transplanted with MLL-AF9 leukemic cells, suggesting that a leukemic state may attenuate the supportive function of *Lepr*+ stromal cells for normal hematopoiesis [24].

The relevance of these proposed concepts and mechanisms for human disease have, however, remained uncertain, primarily since we lack comprehensive insights on LEPR^+^ BMSCs in the human leukemic marrow and their potential pathological interaction with hematopoietic elements.

Here, we performed comprehensive, paired, single cell transcriptional sequencing and networking analyses of all cells in human normal bone marrow (NBM) and AML aspirates.

The data establish a comprehensive taxonomy of the predicted cellular interactions between LEPR^+^ stromal niches, HSPCs and adaptive and innate immune cells in the human NBM and the disruption of these signaling pathways defining AML. Induced deterioration of stromal niche support in AML through inflammatory activation of LEPR^+^ cells is predicted to broadly affect tissue signaling and induce repression of normal hematopoiesis via disruption of signaling towards high-output HSCs and committed progenitors, but is ultimately associated with a favorable prognosis. The data provide human disease relevance to previous findings in model systems and provide an unprecedented resource of intercellular signaling in the human bone marrow defining native and neoplastic hematopoiesis, with a particular emphasis on the communication between HSPCs and their stromal niches.

## RESULTS

### A cellular taxonomy of the human normal bone marrow

To generate a cellular taxonomy of the human NBM, representing both rare hematopoietic stem/progenitor cells (HSPCs) and stromal niche populations, allowing assessment of their cellular diversity and predicted intercellular signaling, we performed single cell RNA sequencing (scRNAseq) on viably frozen bone marrow (BM) aspirates from four healthy donors for allogeneic transplantation (median age 51, range 47-53 years) (Table S1). To ensure robust representation of all BM cell types in the dataset (enabling accurate assessment of population heterogeneity at high resolution), we used flow cytometry to sort and purify all cells in the aspirates into five fractions: the non-hematopoietic stromal (CD45^-^CD235a^-^CD71^-^CD31^-^) and non-stromal/endothelial (CD45^-^ CD235a^-^CD71^-^CD31^+^) fraction, the HSPC fraction (CD45^+^CD34^+^), the myeloid fraction (CD45^+^CD34^-^CD117^+^/CD33^+^), and the non-myeloid (lymphoid) fraction (CD45^+^CD34^-^ CD117^-^CD33^-^) (Fig. S1a). Fractions were pooled into two fractions (HSPC with myeloid and non-hematopoietic with non-myeloid) and subjected to scRNAseq in separate runs and the data of the two separate runs was subsequently merged into a single scRNAseq dataset (Fig. S1a).

We acquired high quality data from 46,740 NBM cells. The purification strategy resulted in robust representation of low-frequency cell populations such as non-hematopoietic stromal cells and hematopoietic stem cells (HSCs) in the dataset (Fig. 1a and S2a).

**Figure 1.**
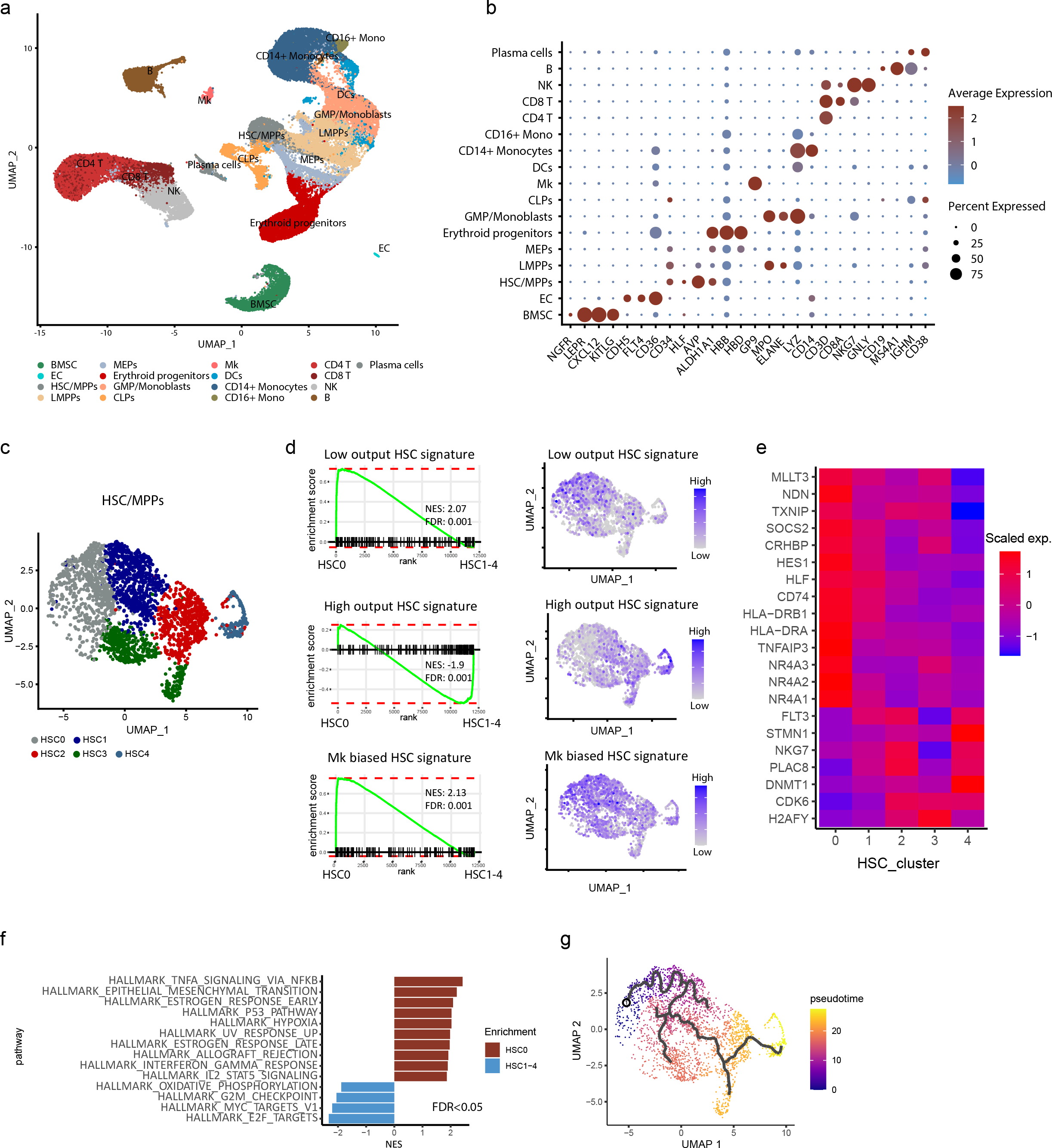
A cellular taxonomy of the human normal bone marrow. (a) Uniform Manifold Approximation and Projection (UMAP) plot of mononuclear cells from bone marrow aspirates, representing 46,740 cells from 4 healthy donors. (b) Expression of cell-type defining genes across all cell types. Color scale and dot size reflect levels and percentages of cells with detectable gene expression. (c) Heterogeneity of the HSC/MPP population reflected in a UMAP plot, representing 2763 cells. (d-e) Identification of low-output HSCs (cluster 0) and high-output (cluster1-4) HSC/MPPs, based on transcriptional homology with these subsets identified in mice. (d) Gene signatures for low-, high-output and megakaryocyte(Mk)-biased HSCs in the HSC/MPP subpopulations. (e) Heatmap showing differential expression of genes between clusters. (f) Differentially expressed transcriptional programs in HSC cluster-0 in comparison to cluster-1-4, as demonstrated by Hallmark analysis. Positive NES (Normalized Enrichment Score) reflects programs enriched in cluster-0, while negative scores indicate enrichment in cluster-1-4. FDR<0.05. (g) Predicted HSC lineage progression by trajectory analysis. Calculated pseudotime is represented by color scale. HSC cluster-0 is set as starting point.

Using Clustifyr, a package to classify cells from scRNAseq data using external references [25], the sequenced cells were classified into bone marrow stromal cells (BMSCs), endothelial cells (ECs), CD34^+^ HSPCs, erythroid progenitors, megakaryocytes (MKs), dendritic cells (DCs), CD14^+^ monocytes (CD14+ Mono), CD16^+^ monocytes (CD16+ Mono), natural killer cells (NKs), CD4^+^ T cells (CD4 T), CD8^+^ T cells (CD8 T), B cells (B) and plasma cells (Fig. S1b). Accuracy of the unsupervised cell annotation was validated by checking the expression of canonical markers for each cell type (Fig. 1b, S1c-d). The BMSC cluster, for instance, specifically expressed BMSC markers *NGFR* (CD271), *PRRX1*, the perivascular MSC marker *LEPR* and the HSPC niche factors *CXCL12* and *KITLG* (Fig. 1b, S1c-d). The HSPC cluster expressed *CD34*, *AVP* and *KIT* (Fig. 1b, S1c-d) and the CD4^+^ T, CD8^+^ T and NK clusters had enhanced expression of *CD3D*, *CD8B* and *NKG7,* respectively (Fig. 1b, S1c-d).

To dissect the cellular heterogeneity within the HSPC fraction, we further classified this subset using the K Nearest Neighbor (KNN) classification method, resulting in distinct subtypes which were subsequently clustered into HSC/MPPs, lympho-myeloid primed progenitors (LMPPs), megakaryocyte and erythroid progenitors (MEPs), erythroid progenitors, common lymphoid progenitors (CLPs) and granulocyte-monocyte progenitors (GMPs)/monoblasts (enriched) subpopulations (Fig. S2a). Annotation of each HSPC subpopulation was based on comprehensive transcriptional analysis including *CD34* expression (highly expressed in HSC/MPPs, LMPPs, CLPs but gradually deceased in GMP/monoblasts and erythroid progenitors) (Fig. S2b), genes indicative of cell cycle status (cells in the HSC cluster were predominantly retained in a non-cycling (G1) phase) (Fig. S2b), gene enrichment scores to identify HSPC subsets [26] (Fig. S2c) and expression of genes indicative of HSPC type (*HLF*, *AVP* and *HES1* for HSCs, myeloid markers *LYZ*, *MPO* and *S100A8/A9* for GMPs, B cell lineage marker *CD79A* for CLPs and erythroid markers *CA1*, *HBB* and *HB* for MEPs) (Fig. S2d-e) [27]. Trajectory analysis was congruent with the view that HSCs are at the basis of a cellular hierarchy giving rise to erythroid, myeloid and lymphoid progeny (Fig. S2f).

### HSC heterogeneity in the human native bone marrow

ScRNA sequencing studies in mice have revealed transcriptional differences amongst HSCs, providing an explanation for their functional heterogeneity. Distinct transcriptomes discern long-term (LT), low-output HSCs that self-renew rather than proliferate or differentiate, from short-term (ST), high output HSCs that provide lineage output towards mature progeny during normal steady-state hematopoiesis [28–30]. Whether a similar hierarchy of HSCs, reflected in their transcriptome, exists within the HSC pool in human native hematopoiesis has remained largely unknown.

Subclustering of the human HSC/MPP population distinguished 5 subsets (HSC 0-4) (Fig. 1c). Cluster 0 constitutes 23.85 +/- 1.18 % of the HSC pool (Fig. 1c). The transcriptional signature of this cluster 0 had strong relationship with the transcriptional signature previously reported for low-output LT-HSCs in mice [28, 31], while clusters 1- 4 displayed strong transcriptional congruence with signatures of high-output ST-HSCs (Fig. 1d).

The ‘low-output’ HSC cluster 0 was also enriched for HSC signatures previously associated with platelet/megakaryocyte-biased HSCs [28, 31], previously demonstrated to reside at the apex of the hematopoietic stem cell hierarchy [32] (Fig. 1d).

Transcriptional similarity of cells in cluster 0 with murine LT-HSCs included expression of genes associated with self-renewal and quiescence such as *Mllt3*, *Socs2*, *Txnip* and *Ndn*, MHC class II components (*Cd74, H2-Eb1(HLA-DRB1)*), and transcription regulators (*Hlf*, *Hes1*) (Fig. 1e and Table S2). Significantly differentially expressed genes in clusters 1-4 included *CDK6*, previously reported to be a critical regulator for the transition of LT-HSCs to ST-HSCs and MPPs [33, 34] and *FLT3* (Fig. 1e).

GSEA demonstrated transcriptional activation of interferon and STAT activation-related inflammatory signaling (TNF signaling via NFκB; Interferon gamma response; IL2- STAT5 signaling) in the HSC-0 cluster (Fig. 1f), reminiscent of an inflammatory transcriptional signature defining a subset of CD74^high^ MHCII^high^ HSCs resistant to myeloablation in mice [35]. Interestingly, human cluster-0 HSCs display concomitant overexpression of genes encoding proteins inhibiting inflammation, including *TNFAIP3* (A20) [36] and all members of the NR4A subfamily of nuclear receptors (*NR4A1*, *NR4A2* and *NR4A3*) (Fig. 1e), previously shown to protect LT-HSCs against DNA damage and repress the proliferative inflammatory response of HSCs [37, 38], while cell cycle related Hallmark gene signatures (E2F targets and G2M checkpoint) were significantly enriched in clusters 1-4. Cell trajectory analysis was congruent with the notion that the HSC-0 subset may be at the basis of a cellular hierarchy with differentiation trajectories towards HSC clusters 1-4 (Fig. 1g).

Collectively, the data reveal the existence of distinct transcriptional subsets within the HSC/MPP population in human native hematopoiesis, suggesting a linear hierarchy from low-output LT-HSCs to high-output ST-HSC and MPPs, that has previously been demonstrated in mice.

### LEPR^+^ stromal cells are the main predicted source of cellular signaling in the human bone marrow

The generation of a comprehensive cellular taxonomy allows assessment of predicted cellular interactions in the human bone marrow. We analyzed the predicted intercellular communication between all retrieved cell types based on possible ligand-receptor crosstalk using CellChat, which predicts the major signaling routes and how they integrate into cellular function, using network analysis and pattern recognition approaches [39], revealing a complicated cell interaction network in which all cell types were involved (Fig. 2a).

**Figure 2.**
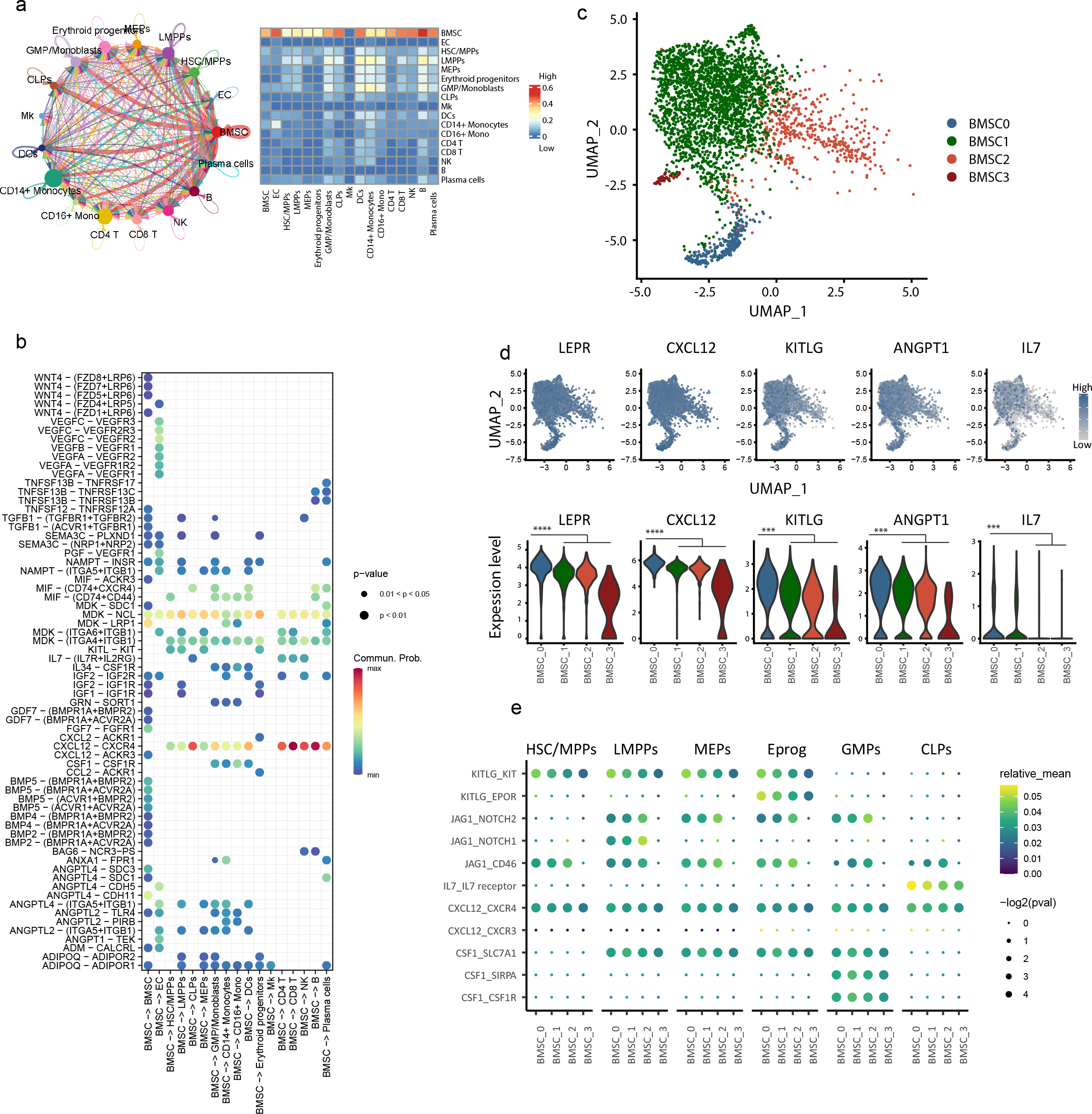
Transcriptional identification of BMSC heterogeneity in the human normal bone marrow. (a) Predicted cellular interactions based on transcriptional networking, identifying BMSCs as the dominant source of signaling to all other cells. In the circle plot, colors represent signal senders and width represents signal strength. In the heatmap, signal strength is represented by the color scale. (b) Predicted ligand-receptor interactions between BMSCs and other cell types. Color scale and dot size represent probability and p-value of interactions, respectively. (c) Heterogeneity of the BMSC population reflected in the UMAP plot representing 3236 cells. (d) Differential expression of *LEPR* and genes encoding key HSPC regulatory factors in BMSC subset-0 represented by UMAP and Violin plots. ***, FDR-adjusted P value (padj)<0.001. ****, padj<0.0001. Differential gene expression analysis is performed using the pseudoDE R package at sample level (pair-wise comparison in individual samples). (e) Relative strength of predicted HSPC-supportive signaling originating from distinct BMSC subsets, as assessed by CellphoneDB. Color scale and dot size represent relative mean strength and p-value of interactions, respectively .

BMSCs were identified as a key ‘communication hub’ in these analyses, with predicted signaling routes to almost all other cell types in the human marrow, with a predicted magnitude of signaling interaction exceeding that of any other cell population in the marrow (Fig. 2a). This predicted signaling included well established interactions of BMSCs with HSPCs via KITLG (encoding stem cell factor, SCF) (to HSC/MPPs, LMPPs, MEPs and erythroid progenitors) [9] and all immune subsets via CXCL12 [10, 11, 40], lymphoid cells via IL7 (to CLPs, CD4^+^ T, CD8^+^ T and NK) [4, 41] myeloid cells via CSF1 and IL34 (to GMP/monoblasts, DCs, CD14+ monocytes and CD16+ monocytes) [42], endothelial cells via VEGF and ANGPTL [43], and predicted autocrine signaling via IGF, BMP, TGFβ and adiponectin, previously implicated in fate-decision signaling (Fig. 2b).

Collectively, the data describe a cellular taxonomy of the stromal and immune environment of the human NBM and their predicted intercellular signaling, unveiling a predicted central role of BMSCs, not only in the homeostatic regulation of HSCs and HSPC subsets, but also all innate and adaptive immune cells, in line with experimental data from mouse studies (Tokoyoda et al. 2010; Comazzetto et al. 2019).

### Stromal cell heterogeneity in the human bone marrow

BMSCs, as shown in mice, represent a heterogeneous population with pleiotropic function in tissue homeostasis and support of distinct hematopoietic cell types [44, 45]. Heterogeneity is, in part, related to anatomical localization (endosteal vs. medullary localized BMSCs), but also within the medullary fraction of BMSCs (likely best represented in bone marrow aspirates), heterogeneity exists in mice [2]. Whether such heterogeneity exists in humans has remained largely unknown. The robust representation of stromal cells in our datasets allowed for addressing this question.

Subclustering of the BMSC population distinguished 4 subsets (BMSC-0-3) (Fig. 2c). All BMSC clusters expressed the sinusoidal stromal HSPC niche marker *LEPR* [9], albeit at different levels, as well as pre-adipocytic markers (*LPL*, *ADIPOQ* and *CD36*) and HSC niche factors (*CXCL12*, *KITLG* and *ANGPT1*) (Fig. 2d, S3a), while expression of osteolineage differentiation markers (*BGLAP, RUNX2, SPP1*), chondrocyte differentiation makers (*SOX9*, *ACAN*, *COL2A1*), fibroblast markers (*S100A4*, *SEMA3C*), and pericyte markers (*NES*, *NG2* and *ACTA2*) could not be detected or at a low level (Fig. S3a).

These findings seem consistent with the notion that the BMSCs captured in our analyses represent the human equivalent of the sinusoidal *LEPR* expressing stromal niche cells in mice that display adipogenic differentiation capacity that have been characterized in mice as adipo-primed BMSCs [24, 46] and are essential for HSPC maintenance [4, 9]. Other microenvironmental cells, such as endothelial cells, pericytes and osteoblasts could not be retrieved from human bone marrow aspirates, or in numbers too low to perform robust analyses (in the case of endothelial cells).

The BMSC-0 subset, comprising 10.5%±5.11% of all stromal cells, displayed the highest expression of *LEPR* and genes encoding critical HSPC regulatory factors (*CXCL12*, *KITLG*, *ANGPT1* and *IL7*) in comparison with other clusters (Fig. 2d). Transcriptome-wide comparison of this subset with the other stromal subsets revealed differential expression of 443 genes (144 upregulated and 299 downregulated, padj<0.05) (Fig. S3b, Table S3).

Ligand-receptor analysis revealed that the BMSC-0 subset had the strongest predicted KITLG-KIT, CXCL12-CXCR4 and IL7-IL7 receptor interaction with HSPCs, while these interactions were less strong in other clusters (BMSC-1, -2 and -3) (Fig. 2e). BMSC-0 derived KITLG was predicted to signal to all HSC subsets (HSC0-4), while other interactions, in particular signaling via the CD74 receptor, was predicted to be strongest to the LT-HSC subset of HSCs (Fig. S3c).

Taken together, the data indicate that heterogeneity exists within the human BMSC population with a minor subset, characterized by the highest expression of genes encoding critical HSPC maintaining factors, predicted to have the strongest interaction with HSPC subsets.

### A cellular taxonomy of human *NPM1* mutated AML

Earlier approaches to establish cellular hierarchies in human AML have focused on hematopoietic cells [47], precluding assessment of interactions between rare cell populations such as HSPCs and their stromal niches and its potential involvement in AML pathogenesis. We, thus, generated scRNAseq tissue maps to establish the cellular taxonomy of the bone marrow in AML with mutations in the gene encoding nucleophosmin (*NPM1*), among the most frequently mutated gene in AML, representing approximately 30% of newly diagnosed AML patients [48].

Bone marrow aspirates from six *NPM1* mutant (*NPM1m*) AML patients (median age 53.5, range 39-59 years) (Table S1) were sorted and subjected to scRNAseq, according to the strategy described for NBM, resulting in transcriptomes of 49,758 cells comprising the taxonomy of the bone marrow in *NPM1*m AML (Fig. 3a). The cells representing the clusters annotated within the NBM could largely be retrieved from the AML bone marrow, although disruption of architecture within defined populations was observed and will be described in the next chapters.

**Figure 3.**
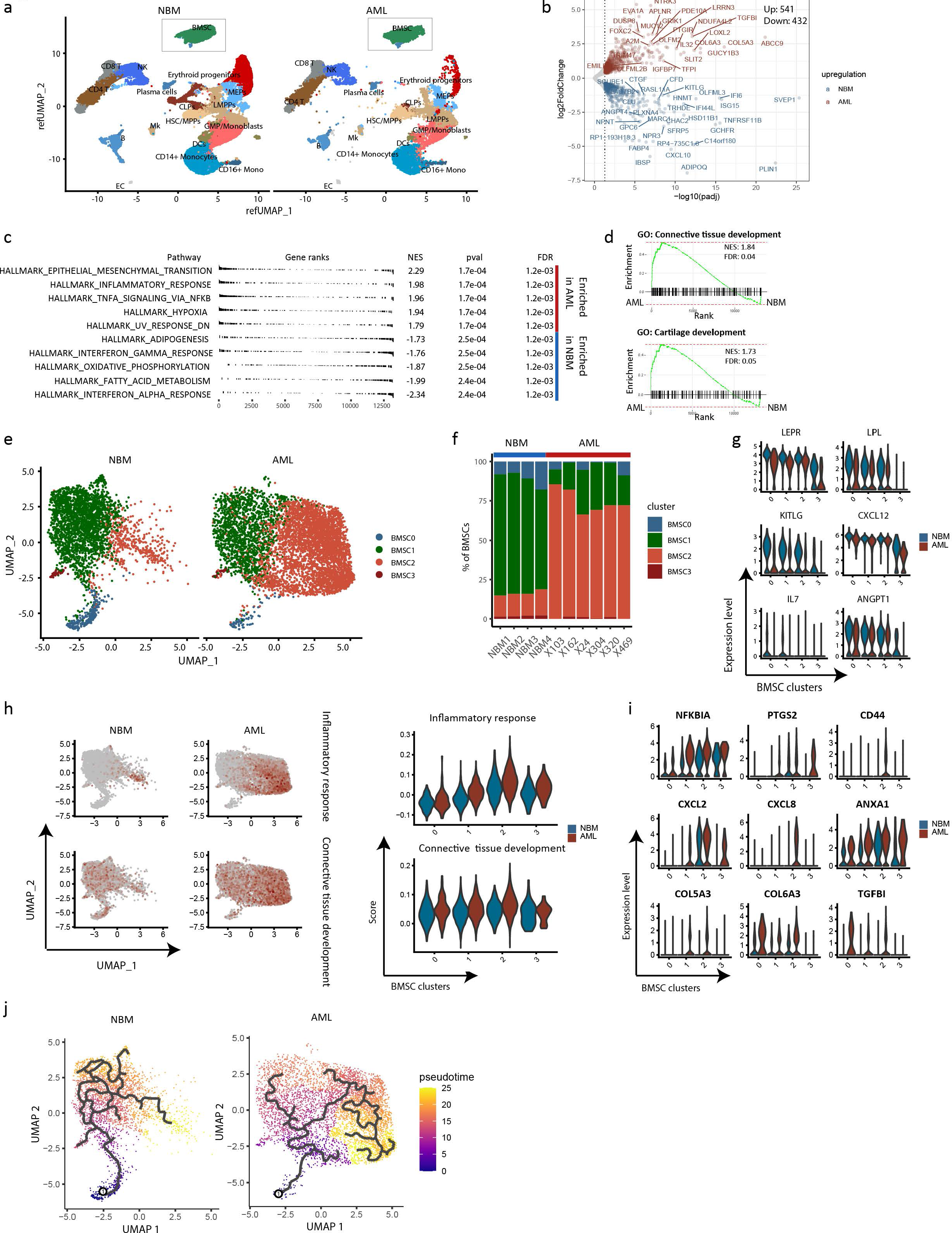
Remodeling of bone marrow stromal cells in *NPM1*m AML. (a) UMAP distribution of bone marrow cells in NBM and AML. For AML, 49,758 cells from 6 patients are presented. (b) Volcano plot of differentially (p-adjusted <0.05) expressed genes in BMSCs in AML versus NBM. Differential expression gene analysis is performed at sample level using the pseudoDE R package. (c-d) Differentially expressed transcriptional programs in BMSCs from AML in comparison to NBM, as demonstrated by Hallmark analysis (c) and GO term (d). Positive NES (Normalized Enrichment Score) reflects programs enriched in AML, while negative scores indicate enrichment in NBM. (e-f) Distribution and frequencies of BMSC subsets in AML and NBM. Note the relative reduction in BMSC cluster-0 and cluster-1, and increase of cluster-2 in AML. (g) Reduced expression of genes encoding HSPC regulatory factors in BMSC subsets in AML. (h-i) Activation of transcriptional programs related to Inflammation and Connective Tissue Development in BMSC cluster-2, as demonstrated by gene signature calculation (h) and expression of inflammation- and ECM remodeling-associated genes (h). (j) Trajectory analysis suggesting inflammatory activation of LEPR+ stromal cells in AML.

### Inflammatory remodeling of LEPR^+^ BMSCs in NPM1m AML

The predicted central role of LEPR^+^ BMSCs as intercellular signal trafficker in the human NBM, raised the question whether, and how, LEPR^+^ BMSC characteristics are disrupted in AML and if alterations could contribute to disease pathogenesis.

Direct comparison of the BMSC transcriptome at population level (outlined in Fig. 3a) in AML to normal controls showed significant (padj< 0.05) differential expression of 973 genes (541 genes upregulated and 432 genes downregulated) (Fig. 3b and Table S4). Among the most upregulated genes in AML were genes associated with inflammatory activation, including *NFKBIA, PTGS2, CD44, CXCL2, CXCL8* and *ANXA1*, and genes associated with extracellular matrix remodeling, including *LOLX2*, *TGFBI*, *COL5A3* and *COL6A3* (Fig. 3b, S4a-S4b), with significant downregulation of genes encoding critical HSPC regulatory genes such as *KITLG*, *CXCL12*, and *IL7* and the gene encoding the HSPC niche marker *LEPR* and pre-adipocyte marker *LPL* (Fig. 3b, S4a-S4b). This was reflected in gene signatures (GSEA Hallmark) consistent with inflammatory signaling (TNFα Signaling Via NFκB; IL2-STAT Signaling; Inflammatory Response) among the top upregulated transcriptional signatures, and downregulation of signatures associated with adipogenesis in AML BMSCs (Fig. 3c, Fig. S4c). Disruption of metabolic pathways was suggested by gene signatures indicative of glycolysis in AML BMSCs (Hallmark Hypoxia; Hallmark Glycolysis) versus oxidative phosphorylation and fatty acid metabolism in normal BMSCs (Fig. 3c, Table S4). In addition, the GO terms Connective Tissue Development and Cartilage Development were also significantly increased in the AML BMSCs (Fig. 3d, Fig. S4c), indicative of remodeling of the extracellular matrix.

To assess how this disruption of transcriptional programs in the overall BMSC population relates to BMSC heterogeneity in AML, we performed subclustering of the BMSC population (Fig. 3e). This revealed near loss of the cluster predicted to have the strongest interaction with HSPCs (BMSC-0) (3,53%±4.84% vs. 11,02%±3,39% in AML and NBM respectively; p=0.044), with a concomitant relative increase of BMSC-2 (46.4%±6.81% vs. 7.85%±1.24%; p=0.009) (Fig. 3e-f, S4d). Levels of genes encoding critical HSPC maintaining factors were significantly reduced in all BMSC subsets in AML, which was most pronounced for *KITLG* and *IL7*, with dramatic impaired expression in BMSC-0 (Fig. 3g).

The BMSC-2 cluster in AML was transcriptionally characterized by upregulation of genes and transcriptional programs (Hallmark: Inflammatory Response) associated with inflammation and genes and signatures related to ECM remodeling (Fig. 3h-3i). This pattern of inflammatory disruption of BMSC architecture and reduced expression of HSPC factors was consistent among all six *NPM1*m AML samples examined (Fig. S4a- d).

Stromal inflammation was confirmed *in situ* by demonstrating increased CD44 protein expression (which is a marker of stromal activation and inflammation [49] in CD271^+^CXCL12^+^ BMSCs in NPM1m AML (by immunofluorescence on bone marrow biopsies) (Fig. S4e-f).

Trajectory analysis suggested that the relative reduction of BMSC-0 and increase in the inflammatory BMSC-2 subset may reflect a linear hierarchy characterized by a gradual increase in inflammatory activation accompanied by a loss of expression of HSPC regulatory factors (Fig. 3j).

Taken together, the data are congruent with the view that in *NPM1*m AML, BMSCs are remodeled by inflammatory activation, resulting in dramatic expansion of an inflammatory subset with a concomitant loss of the BMSC subset predicted to support the maintenance of normal HSPCs.

### Inflammatory remodeling of stromal niches is predicted to repress high-output HSCs and committed progenitors in AML with relative resistance of LT-HSCs and clonal cells to the deprivation of niche factors

Next, we sought to explore the consequences of BMSC remodeling in AML for the residual normal and leukemic hematopoiesis. AML is characterized by suppression of normal hematopoiesis and expansion of clonal cells, but the cellular and molecular mechanisms promoting these processes have remained incompletely understood. Mouse modeling has suggested important contributions of stromal HSPC niches, but the relevance for human AML has remained uncertain.

The localization of *NPM1* mutations at the 3’ terminal coding region of the gene, captured by polyA-RNAseq, allowed us to distinguish mutated, clonal AML cells from residual normal HSPCs by assessing the mutational state of all cells in the dataset (Fig. 4a and section methods). Analysis of mutation status confirmed that adaptive immune subsets (CD4/CD8 T-cells, plasma cells and B-cells), NK cells and BMSCs are of non-clonal origin in AML. The very small fraction of *NPM1m* cells within these fractions differentially and highly expressed myeloid genes (*AZU1*, *LYZ*, *MPO*, and *ELANE)* (data not shown), indicative of blast contamination. Mutated cells reside predominantly within the LMPP and GMP subsets of the HSPC fraction (Fig. 4a-b).

**Figure 4.**
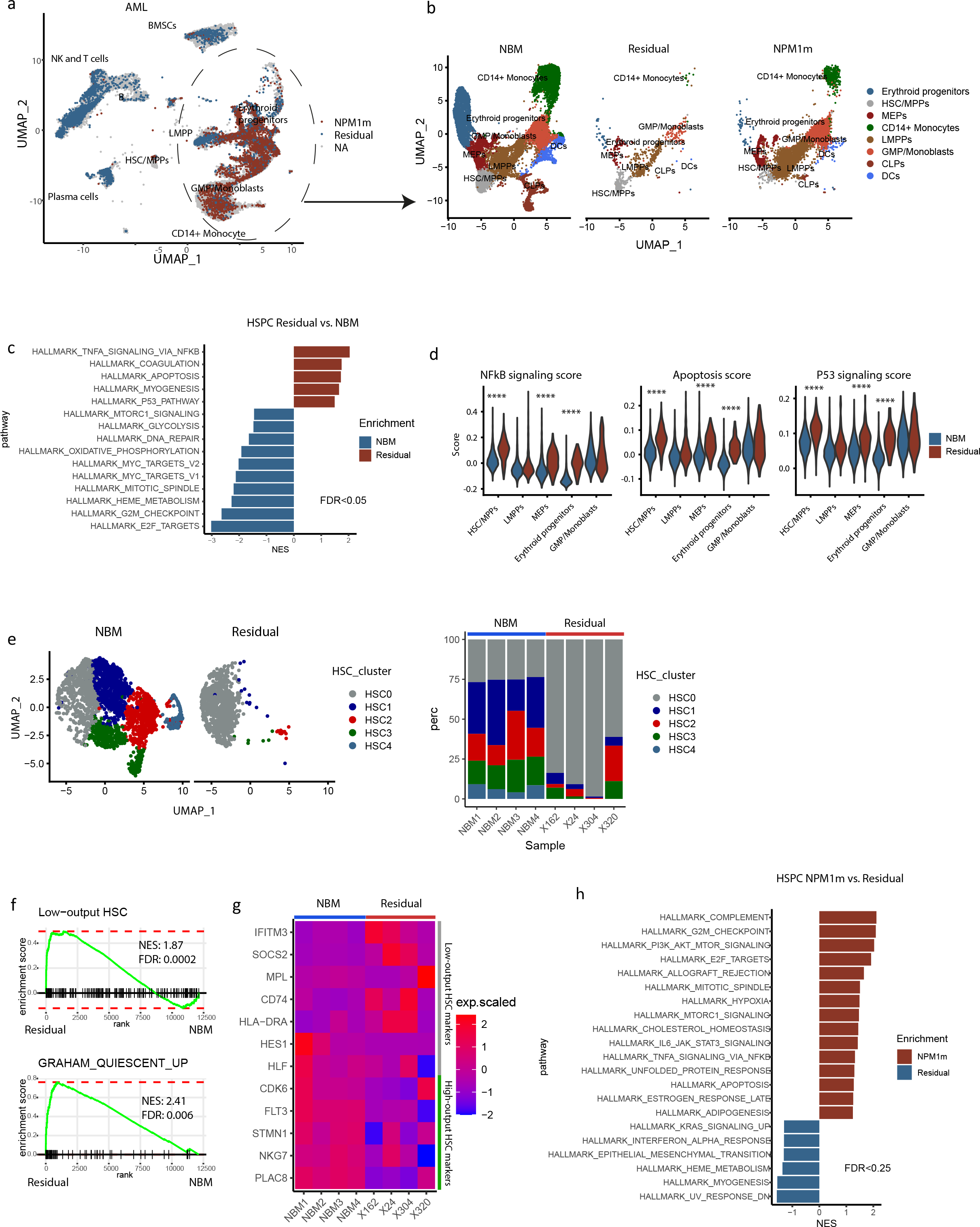
Normal hematopoiesis is suppressed in AML patients, while NPM1 mutant cells display relative resistance. (a) UMAP distribution of *NPM1* mutant (*NPM1m*) (red) and residual normal cells (blue) in the AML bone marrow. NA (grey), not assignable. (b) Distribution of normal, residual normal and *NPM1m* cells within the HSPC and myeloid fractions in NBM and AML. Note that in AML, the majority of cells in the HSC/MPPs fraction is of residual normal origin, while the vast majority of cells within the LMPP and GMP fractions carry the leukemic mutation. (c) Differentially expressed transcriptional programs in residual normal HSPCs in AML compared to HSPCs in NBM, as demonstrated by Hallmark analysis. Positive NES (Normalized Enrichment Score) reflects programs enriched in residual normal, while negative scores indicate enrichment in NBM. All cells in HSC/MPPs, LMPPs, MEPs, Erythroid progenitors and GMP/Monoblasts clusters were analyzed. (d) Overexpression of hallmark transcriptional signatures indicative of NFkB signaling, Apoptosis and P53 signaling in residual normal HSPC in comparison to their counterparts in the human NBM . ****, padj<0.0001 by Wilcoxon test. (e) Distribution and frequencies of HSC/MPP subsets of residual normal HSPCs in comparison to NBM. Note the relative depletion of the high-output HSC/MPP clusters-1-4 with relative preservation of transcriptional low-output HSCs in AML. In two AML samples insufficient cells could be retrieved in the HSC/MPP subset for analysis. (f) Enrichment of transcriptional programs indicative of low-output, quiescent, HSCs in residual normal HSC/MPPs in AML in comparison to their counterparts in the human NBM. (g) Heatmap for low- and high-output HSC marker genes in NBM and residual normal HSC/MPP population across all samples. (h) Differentially expressed transcriptional programs in *NPMm* cells within the HSPC subsets in comparison to residual normal HSPCs, as demonstrated by Hallmark analysis. Positive NES (Normalized Enrichment Score) reflects programs enriched in *NPM1m* HSPCs, while negative scores indicate enrichment in residual normal HSPCs. All cells in HSC/MPPs, LMPPs, MEPs, Erythroid progenitors and GMP/Monoblasts clusters were used in the analysis.

The remodeling of stromal niches in human AML, in particular the loss of stromal subsets predicted to have the strongest supportive interaction with HSPCs, and the marked downregulation of KITLG encoding stem cell factor (SCF), is predicted to have major consequences for residual normal hematopoiesis. Depletion of Scf from *Lepr*+ BMSCs in mice results in cytopenia, hypocellularity of the marrow and a reduction in HSC number[5], indicating that stromal SCF is critical for the maintenance of normal HSCs and hematopoiesis. *Lepr*+ BMSC derived Scf is also critical for the maintenance of kit+ restricted hematopoietic progenitors (in particular CLPs and MEPs and to a lesser extend CMP and GMP) [9].

Ligand-receptor analyses confirmed that BMSC-HSPC interactions deemed critical for normal HSPC maintenance, survival and proliferation (such as KITLG-KIT and CXCL12- CXCR4) were attenuated in residual normal HSPC subsets in AML in comparison to their counterparts in NBM (Fig. S5a). We therefore evaluated the cellular changes in the residual normal HSPC compartment associated with predicted disruption of niche signaling. The HSPC fraction in AML displayed a relative reduction of erythroid progenitors (17.41%±9.77% vs. 6.31%±5.74% of the HSPC fraction in NBM and AML respectively, p=0.1) and GMP/monoblasts (22.08%±3.29% vs. 12.14 %±7.15%, p=0.02), as well as near loss of the CLP population (8.85%±3.79% vs. 0.15%±0.378%, p=0.019) (Fig. 4b), recapitulating observations in mice with depletion of Scf from *Lepr*+ BMSCs.

In line with the notion that residual normal HSPCs in AML may be affected by the loss of stromal niche factors, and in particular SCF propagating survival and proliferation signals, which occur via activation of PI3K-AKT-MTORC signaling [50, 51], GSEA and hallmark pathway analyses demonstrated enrichment of inflammatory and apoptotic signatures in residual normal subsets in comparison to their counterparts in the human NBM (Hallmark: TNFα via NFκB; Hallmark: apoptosis and Hallmark: P53 pathway) with concomitant depression of signatures indicative of MTORC signaling, active cellular metabolism and proliferation (Hallmark: TORC1 signaling; Hallmark: E2F targets; Hallmark: G2M checkpoint, Hallmark: Mitotic spindle and Hallmark: Oxidative phosphorylation) (Fig. 4c,d). Interestingly, these transcriptional programs indicative of cellular stress were activated most extensively in residual normal cells within the HSC/MPP, MEP and erythroid progenitor populations (Fig. 4d), the progenitor cells known to be most repressed in AML (resulting in anemia and thrombocytopenia).

The CD34^high^ HSC compartment was largely of residual normal origin (with only 3.66%±2.88% of cells within this HSC/MPP population detected positive for *NPM1* mutation) (Fig. 4a), in line with the existing notion that *NPM1*m AML has low expression of CD34 and may find its origin in transformation of committed progenitors [52]. Re-clustering of the HSC population showed that, strikingly, the ST, high-output, HSC/MPP subsets (HSC-1-4) were almost completely depleted from the residual normal HSC pool in AML (74.78%±1.29% vs. 16.47%±16.12% in NBM and AML, respectively, p=0.005) with relative conservation of the LT, low-output, HSC-0 subset (Fig. 4e). In line with this observation, the gene signatures indicative of low-output HSCs and quiescence were significantly enriched in the residual HSCs (Fig. 4f), while the high-output HSC marker genes such as *FLT3*, *STMN1*, *PLAC8* and *MPO* were downregulated (Fig. 4g, Table S5). The finding that this specific HSC subset, predicted to be the LT-repopulating subset of HSCs, resistant to chemotherapy by analogy with experimental findings in mice, resists inflammatory stress, seems consistent with the longstanding observation that the human BM can reconstitute normal hematopoiesis after chemotherapeutic eradication of leukemic cells.

Importantly, *NPM1*m cells within the HSPC fraction, predominantly found within the LMPP and GMP subsets (Fig. 4b), showed preservation of transcriptional MTORC1 and survival signaling (Fig. 4h), suggesting relative resistance to the factors driving the cellular stress in comparison to residual normal HSPCs. Moreover, several interactions were predicted between inflammatory molecules encoded by genes differentially expressed in the ‘inflamed’ BMSC subset (BMSC-2) and the LMPP-like, GMP-like and/or monocyte-like *NPM1*m AML cells, including HGF-CD44, IL6-IL6 receptor and JAG1-NOTCH1/NOTCH2 interactions (Fig. S5a), which have been reported to drive the initiation, proliferation and/or chemo-resistance of AML [17, 53, 54].

Collectively, the scRNAseq data, in conjunction with previously established relevance of niche factors for normal hematopoiesis in mice, support the view that niche remodeling by inflammatory activation and suppression of HSPC maintenance factors in AML suppress normal hematopoiesis, while LT-HSCs and *NPM1*m leukemic cells are relatively resistant to the deprivation of supportive signaling from stromal niches. In addition, inflamed BMSCs factors express factors previously associated with leukemia propagation. Niche remodeling is thus predicted to be a driving force in the competitive advantage of mutated cells over their normal counterparts in leukemogenesis.

### Inflammatory remodeling and loss of *KITLG* expression from BMSC niches and the related repression of normal hematopoiesis in AML may be driven by TNF**α** from activated immune cells

The scRNAseq analyses suggested that the disruption of BMSC architecture in AML may reflect inflammatory activation of LEPR^+^ BMSCs. To provide experimental support for this view, we tested whether secreted inflammatory factors could induce the inflammatory alterations observed in the BMSC compartment in AML. The significantly enriched gene signature ‘TNFα Signaling Via NFκB’ (Fig 3c) suggested that TNFα may be one such factor. TNFα levels have earlier been demonstrated to be increased in the plasma of AML patients [55] and *TNF* is overexpressed in AML at the transcriptional level (Fig. S6a). The scRNAseq data showed that *TNF* is highly expressed in adaptive and innate immune cells (CD8 T cells and NK cells) and myeloid lineage cells (GMP/monoblasts and CD14+ monocyte populations) in *NPM1m* AML (Fig. S6b-S6c), while its canonical receptor *TNFRSF1A* is predominantly expressed in LEPR^+^ BMSCs (Fig. S6d), predicting that LEPR^+^ BMSCs could be one of the most affected cell types in AML by TNFα signaling.

To test whether TNFα can induce inflammatory remodeling of LEPR^+^ BMSCs and attenuate normal hematopoiesis, we performed an *in vivo* experiment, injecting C57BL/6 mice intraperitoneally with recombinant TNFα (Fig. 5a). This resulted in cytopenia (anemia and thrombocytopenia) (Fig. 5b) and a significant reduction of the frequency of LEPR^+^ stromal cells within the non-endothelial (CD45^-^Ter119^-^CD31^-^ CD144^-^CD51^+^Sca1^-^) niche (Fig. 5c,d). RNAseq confirmed high levels of *Kitlg* and *Cxcl12* expression, specifically in the LEPR+ stromal cell population, in line with the notion that these cells represent HSPC niches (Fig. 5e) [5]. TNFα exposure induced overexpression of inflammatory markers, including *Cd44* and *Cxcl2*, with concomitant downregulation of *Kitlg* and *Cxcl12* (Fig. 5e), in line with the notion that TNFα leads to stromal inflammatory remodeling with subsequent reduction of HSPC niche cells, recapitulating findings in human AML. This was further associated with a reduction in bone marrow cellularity and depletion of distinct subsets of HSPCs, in particular MPPs (Lin^-^cKIT^+^Sca1^+^CD48^-^CD150^-^), CMPs (Lin^-^cKIT^+^Sca1^-^CD34^+^CD16^-^), GMPs (Lin^-^cKIT^+^Sca1^-^CD34^+^CD16^+^) and MEPs (Lin^-^cKIT^+^Sca1^-^CD34^-^CD16^-^), resembling the reduction in MPPs and committed HPC fractions in human *NPM1*m AML.

**Figure 5.**
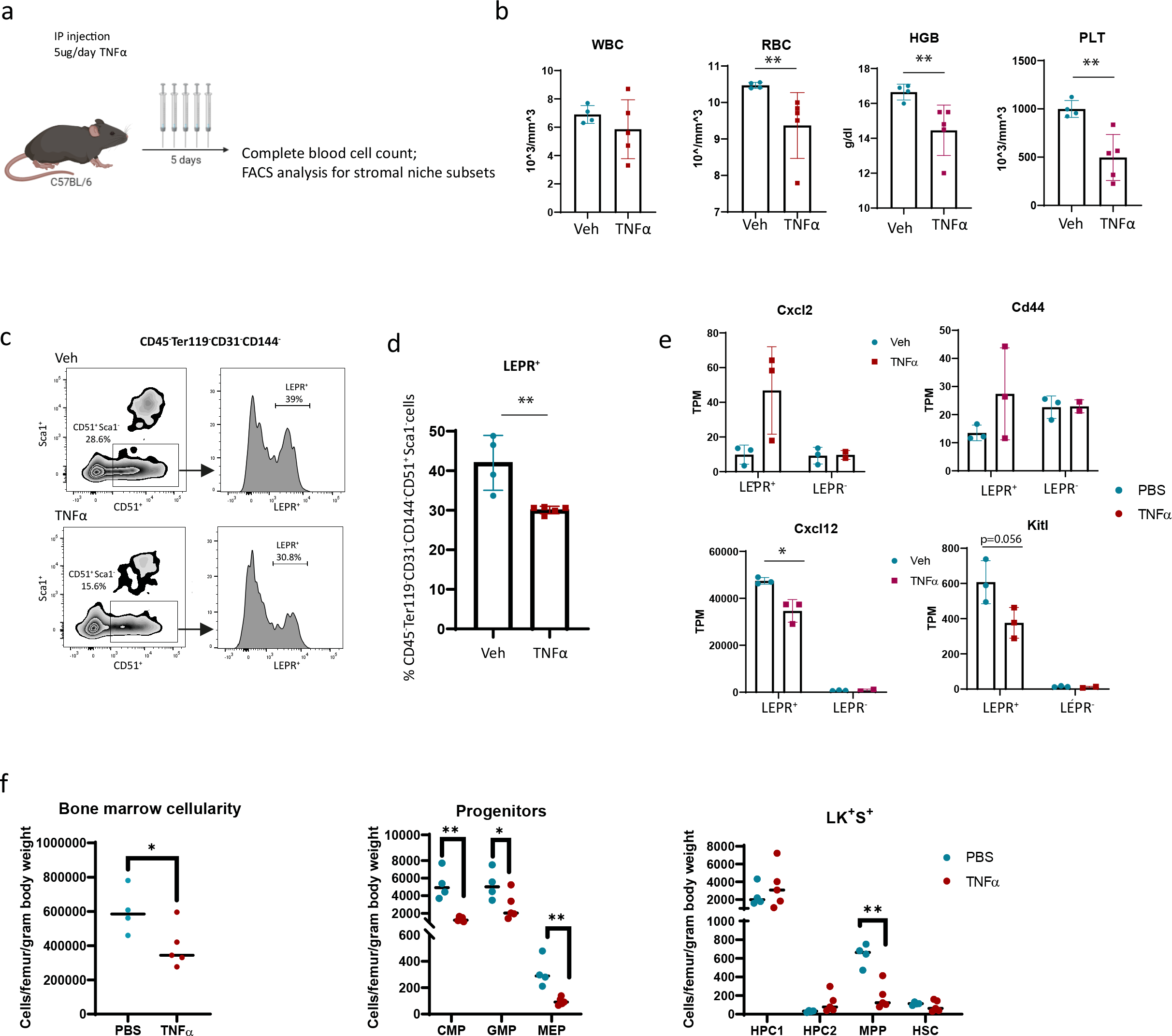
TNF**α** induces inflammatory remodeling of stromal niches and a reduction in HSPCs numbers in mice. (a) Experimental design of TNFα administration to C57BL/6 mice. Five daily i.p. injections at a dose of 5ug were administrated followed by flow-cytrometric assessment of bone marrow stromal niches in collagenased bone fractions. (b) Cytopenia (anemia and thrombocytopenia) in TNFα treated mice. (c-d) Relative loss of LEPR+ BMSCs within the niche fraction upon TNFα exposure. (e) Expression of inflammatory makers and HSPC niche factors in CD51^+^ LEPR^+^ and CD51^+^LEPR^-^ BMSCs after TNFα injection in mice. *, p<0.05 by unpaired t-test. (f) Number of total bone marrow cells, committed progenitors, and LKS HSPC subtypes in mice after TNFα injection. *, p<0.05. **, p<0.01 by unpaired t-test.

To begin testing the functional significance of inflammatory BMSC niche remodeling on normal vs. leukemic hematopoiesis using ex vivo co-culture system, we exposed human HS-5 stromal cells to TNFα (10ng/ml) for 24 hours to trigger the inflammatory response, followed by co-culturing with either human CD34^+^ HSPCs or primary *NPM1*m AML cells for 72 hours (Fig.S6e). This resulted in a significant reduction of immunophenotypic HSPCs (CD45^+^Lin^-^CD34^+^) and CFU-GEMM (Fig. S6e), while the number of *NPM1*m AML cells was not affected by the inflammatory activation of stromal cells induced by TNFα (Fig. S6e). In line with these *in vitro* findings, *in vivo* administration of TNFα in a well-established MLL-AF9 mouse transplantation model of AML did not result in a reduction of leukemic cells (Fig. S6f).

Together, the data fit a model in which overexpression of TNFα, perhaps in part by activation of the innate and adaptive immune system in *NPM1*m AML, results in suppression of hematopoiesis with relative resistance of clonal leukemic cells. This is associated with inflammatory remodeling of stromal niches predicted to affect the survival and proliferation of HSPC subsets, although direct effects of TNFα on HSPCs have also been demonstrated [56] and cannot be excluded.

LEPR^+^ BMSC transcriptional remodeling in AML, conversely, is predicted to affect cellular signaling to lymphoid immune subsets (CD4 T, CD8 T, NK, B and plasma cells) in the leukemic bone marrow (Fig. S5b). This complex cellular taxonomy of the intercellular signaling defining human AML is anticipated to provide a resource to instruct future experimental interrogation of the relevance of these interactions.

### Inflammatory stromal activation and impaired expression of HSPC factors is a biologic commonality in AML, albeit variable between genetic subtypes and risk groups

We next asked the question whether the inflammatory remodeling of BMSCs in AML was restricted to *NPM1*m cases, or rather a biologic commonality in AML across different, distinct, genetic subtypes. To answer this question, bulk RNAseq was conducted on highly purified CD45^-^CD71^-^CD235a^-^CD31^-^CD271^+^ BMSCs isolated from a cohort of 62 AML patients (Fig. S7a), uniformly treated within an intensive chemotherapy clinical trial [57] and selected to represent the mutational landscape of AML (Table S1). Purity of the sorted stromal population was confirmed by excluding expression of hematopoietic transcripts (including CD45 (*PTPRC*), *CD34*, *MPO* and *GYPA*) (Fig. S7b) and expression of canonical stromal markers (CD271 (*NGFR*), *COL1A1*, CD90 (*THY1*) and *PRRX1* (Fig. S7b).

Overall, 1579 genes were differentially expressed in the AML BMSCs in comparison to age-matched normal controls (n=8) (Table S6), including 1413 upregulated and 166 downregulated genes (Table S6). GSEA analyses revealed differentially expressed gene sets, very similar to those identified by scRNAseq in the *NPM1*m subset, including enrichment of the signatures indicative of inflammatory activation (TNFα signaling via NFκB; inflammatory response) and hypoxia, as well as depletion of gene sets indicative of adipogenesis and oxidative phosphorylation) (Fig. 6a). Consistently, expression of the NFκB-related inflammatory genes/cytokines such as *NFKBIA*, *CD44*, *CXCL2, CXCL3*, *CXCL8*, *CCL2*, *LIF* and *PTGS2* were significantly upregulated in AML and adipogenesis markers (*LPL*, *ADIPOQ* and *CD36)* significantly downregulated (Fig. 6b, Table S6). Similarly, expression of *LEPR* and genes encoding the HSPC regulatory factors CXCL12, KITLG and ANGPT1 was significantly reduced in AML BMSCs (Fig. 6b), in line with the scRNAseq data from *NPM1*m patients. Notably, inflammatory activation of BMSC in AML (as represented by the TNFα via NFκB score) was strongly associated with the reduction in expression of HSPC niche genes such as *LEPR*, *CXCL12, KITLG* and *ANGPT1* (Fig. S7c), congruent with the notion that inflammatory activation results in reduction of HSPC niches and downregulation of HSPC factors, as established in our in vivo experiments (Fig. 5c-e).

**Figure 6.**
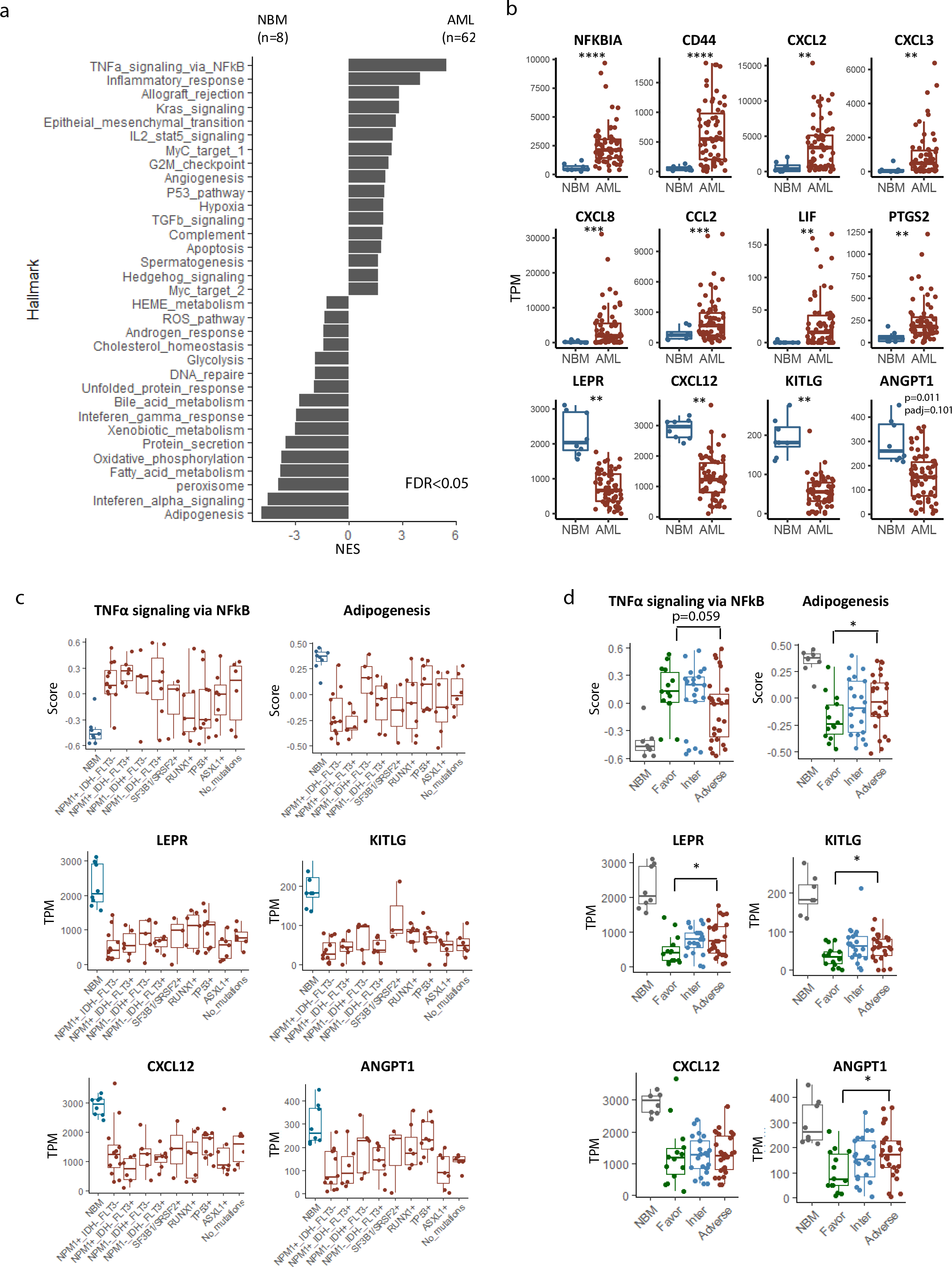
Inflammatory stromal activation and impaired expression of HSPC factors is a biologic commonality in AML, albeit variable between genetic subtypes and risk groups. (a) Differentially expressed gene programs in BMSCs from AML (n=62) in comparison to NBM (n=8) as assessed by Hallmark GSEA. (b) Differential expression of genes encoding inflammatory cytokines/modulators and HSPC niche markers/factors in BMSCs from AML in comparison to NBM. **, padj<0.01, ***, padj<0.0001. TPM = Transcripts Per Million. (c) Enrichment of genesets and expression of genes encoding HSPC niche markers/factors in BMSCs from AML patients related to mutational status. Enrichment scores are calculated using the Gene Set Variation Analysis (GSVA) program. (d) Enrichment of genesets and expression of HSPC niche markers/factors in BMSCs from AML patients related to distinct ELN2017 genetic risk categories. Wilcox test is applied for statistical analysis. *, p-value<0.05. ELN=European Leukemia Network.

To obtain insight into the heterogeneity of these signatures among genetic subtypes, scores were allocated to individual samples and categorized among genetic subtypes using Gene Set Variation Analysis (GSVA) [58] (Fig. 6c), revealing that inflammatory scores were relatively high in the *NPM1*m patients and tended to be lower in patients carrying either a *RUNX1*, *ASXL1* or *TP53* mutation, although heterogeneity existed within these genetic subsets (Fig. 6c). Correlation of BMSC signatures to genetic risk categories as defined by the ELN [59] (shown to be of prognostic relevance in our cohort of 62 patients, as expected) (Fig.S7e), revealed that patients in the favorable risk category displayed the most pronounced NFκB inflammatory activation and suppression of niche supportive factors (*LEPR*, *KITLG* and *ANGPT1*) (Fig. 6d), while BMSCs in the adverse risk group tended to be less perturbed in comparison to NBM.

Together, the data demonstrate that the remodeling of BMSCs, revealed by scRNAseq in *NPM1*m patients and defined by activation of inflammatory signaling and reduced expression of HSPC maintenance factors associated with the loss of the HSPC niche subset, is a biologic commonality in AML, and that the extend of this remodeling may vary within and between genetically defined subsets of patients.

### Stromal inflammatory niche remodeling and associated gene signatures are associated with favorable outcome in AML

We next asked the question whether inflammatory remodeling of stromal niches is related to clinical outcome in AML. The loss of stromal HSPC niche cells has recently been associated with better outcome upon chemotherapeutic treatment in AML in a mouse model [16]. In the MLL-AF9 AML model, genetic depletion of stromal niche cells resulted in delayed relapse after cytarabine treatment [16]. The finding that the HSPC niche (BMSC-0) subset of stromal cells is depleted in AML (to varying degrees) as a result of inflammatory remodeling, prompted us to interrogate the prognostic value of a transcriptional signature reflecting BMSC remodeling in the context of intensive chemotherapy in human AML.

In order to analyze the correlation between BMSC heterogeneity with clinical outcome in AML, we made a ‘by-proxy’ assessment of BMSC heterogeneity from bulk transcriptomes of all 62 patients using CIBEROSRTx, a machine learning method to impute gene expression profiles and provide an estimation of the relative abundance of cellular subsets in a mixed population[60]. The BMSC scRNAseq dataset was used as reference and relative abundance of all 4 BMSC subsets was computationally retrieved (Fig. 7a-b).

**Figure 7.**
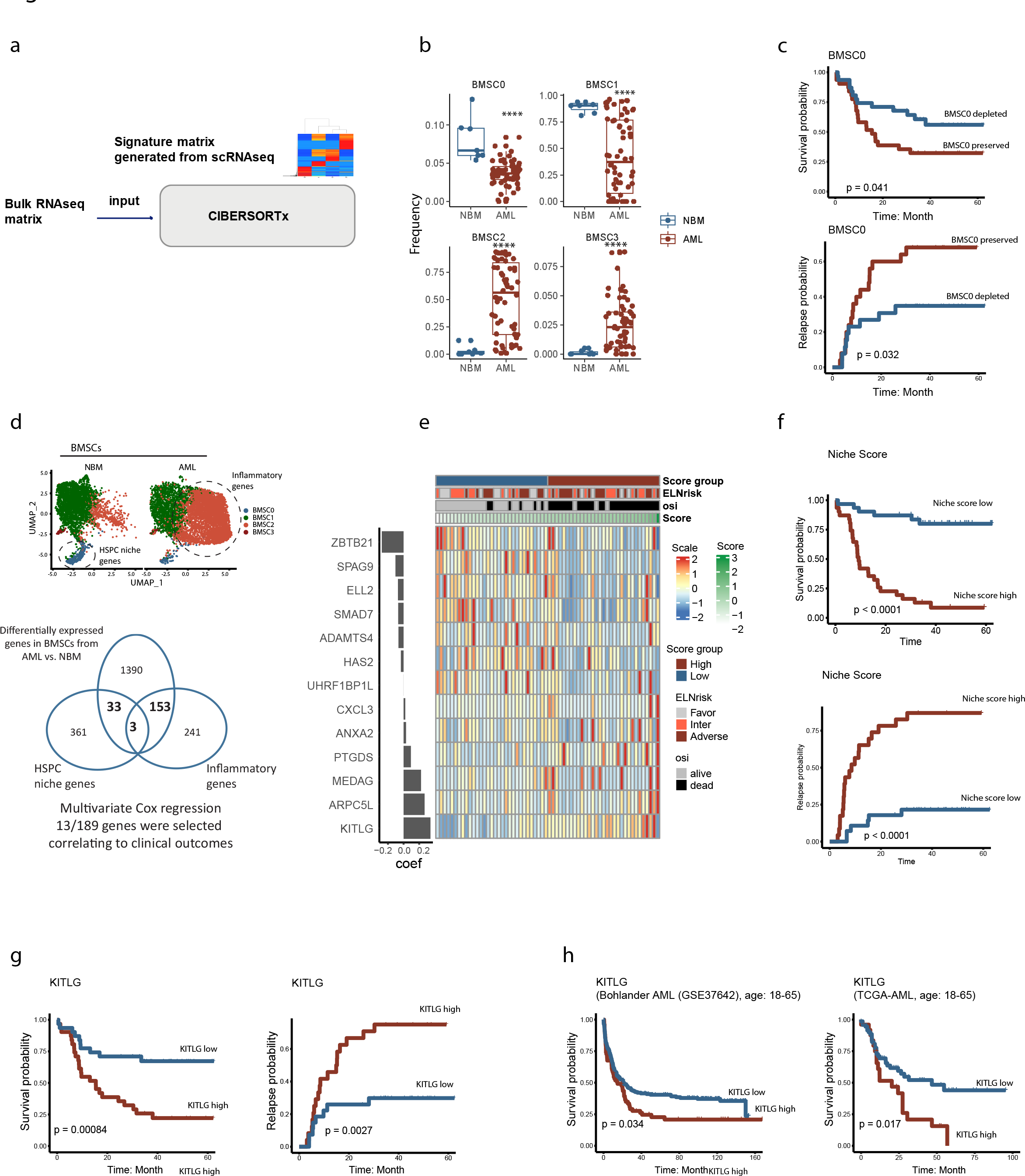
Gene signatures reflective of inflammatory niche remodeling are associated with favorable clinical outcome in AML. (a) Workflow for prediction of BMSC subset representation in BMSC bulk RNAseq data using CIBERSORTx. The BMSC scRNAseq data matrix was used as reference. (b) Predicted BMSC subset size in the larger cohort of AML patients (n=62). Note the variable relative depletion of cluster-0 and cluster-1 in AML with a concomitant relative increase in clusters 2 and 3. ****, p<0.0001. (c) Kaplan-Meier curves for overall survival (OS) and relapse probability indicating improved outcome in patients with the predicted loss of BMSC cluster 0. Cutoff= median frequency. Statistical significance is determined by log-rank test. (d) Construction of a gene signature reflective of niche remodeling (inflammatory activation and relative loss of BMSC cluster 0) in AML. 189 genes at the intersection of the genes differentially expressed in the BMSC-0 vs. other clusters in NBM, the genes differentially expressed in the BMSC-2 vs. other clusters in AML, and the genes differentially expressed in BMSCs in human AML vs. NBM were selected and tested for their correlation with outcome by Multivariate Cox regression survival analysis. 13/189 genes were identified that were associated with outcome. (e) Coefficient plot (left) and heatmap (right) of the 13-gene signature. The Gene Expression Profile score (GEP score) in each patient was calculated using the coefficient and z-scale of the genes. Based on the median GEP score, patients were stratified into two groups, with a high score reflecting relative preservation of niche integrity and a low score reflecting niche inflammatory disruption. (f) Kaplan-Meier curves for overall survival (OS) and relapse probability indicating improved outcome in patients with niche inflammatory disruption. Cutoff= median score. Log-rank test is used for statistical analysis. (g) Kaplan-Meier curves for overall survival (OS) and relapse probability indicating improved outcome in patients with lower *KITLG* expression in BMSCs. Cutoff= median TPM. Log-rank test is used for statistical analysis. (h) Kaplan-Meier curves for overall survival (OS) indicating improved outcome in age-matched AML patients with lower *KITLG* expression in whole bone marrow in Bohlander AML (GSE37642) cohort and TCGA-AML cohort. Cutoff = 75th Percentile normalized counts. Log-rank test is used for statistical analysis.

In line with our scRNAseq observations in *NPM1m* patients, the predicted size of the BMSC-0 subset was significantly reduced in this larger AML population, with a concomitant increase in the size of the inflamed cluster (BMSC-2) (Fig. 7b). Interestingly, stratifying the AML patients into a ‘BMSC-0 preserved’ group and a ‘BMSC-0 depleted’ group based on the median level, revealed that depletion of the BMSC-0 niche subset was significantly associated with better overall survival (5-years OS 57.2 % vs. 31.0% of patients; p=0.041) and reduced risk of relapse (32.0% vs. 65.2% of patients; p=0.032) (Fig. 7c).

Next, we sought to refine this analysis and get better insight into the genes in the remodeled stromal cells that shape this association between niche remodeling and clinical outcome in AML. To this end, we generated a list of genes significantly differentially expressed in BMSCs in human AML (in comparison to BMSCs in NBM) that intersected with inflammation-associated genes (differentially expressed in AML BMSC cluster 2 vs. other clusters in AML) and HSPC niche genes (differentially expressed genes in NBM BMSC-cluster 0 vs. other clusters in NBM), resulting in a list of 189 genes (Fig. 7d, Table S7).

By constructing a penalized multivariable Cox-regression model through nested cross-validation [61], 13/189 genes were identified to be correlated to overall survival (OS) (Fig. 7e). The strongest negative association (coefficient: 0.34; overexpressed in the HSPC niche population and related to poor prognosis) was found for *KITLG,* indicative of a positive correlation between the loss of HSPC niche population and favorable outcome. Other genes overexpressed in HSPC niche population (BMSC-0) and associated with poor OS include *ARPC5L*, encoding an actin related protein involved in cell migration, *MEDAG*, a positive regulator for adipocyte differentiation, and *PTGDS*, encoding a glutathione-independent prostaglandin D synthase that catalyzes the conversion of prostaglandin H2 (PGH2) to prostaglandin D2 (PGD2). Upregulation of the inflamed cluster (BMSC-2) genes *ZBTB21, SPAG9* and *ELL2*, on the other hand, were strongly associated with a favorable outcome.

Based on this 13 gene signature, a score was calculated that allowed for stratifying the AML patients into a ‘BMSC-inflammatory remodeling’ group (score^low^) and a ‘BMSC niche-preserved’ group (score^high^) based on the median level (Fig. 7e). The low score, ‘BMSC-inflammatory remodeling’ group had significantly better OS (5-years OS 80% vs. 8.6%; p<0.0001) and lower relapse probability (5-years relapse probability 13% vs. 78.4%; p<0.0001) (Fig. 7f). Importantly, this significant survival benefit was found throughout all distinct ELN2017 genetic risk categories (Fig. S8a).

Finally, we sought to confirm our finding that stromal niche factors, associated with inflammatory remodeling, are associated with outcome in an independent cohort of AML patients. While many of the factors in the prognostic stromal niche gene signature were either not specifically expressed in stromal cells or not detected in published transcriptional datasets of AML, we found the gene *KITLG* to be 1. an important determinant of the prognostic value of the stromal prognostic gene signature (Fig. 7e, 7g), 2. to be specifically expressed in BMSCs in the AML taxonomy (Fig. S8b) and 3. to be detectable in publicly available transcriptional datasets generated from bone marrow aspirates of AML patients (18-65 years, treated with intensive chemotherapy), namely the TCGA-LAML [62] and Bohlander (GSE37642) [63] cohorts, making it the ideal candidate gene to confirm our findings in independent cohorts. In these datasets, stratifying AML patients into a ‘*KITLG*-low’ group and a ‘*KITLG*-high’ group, based on the 75^th^ percentile expression (to clearly discriminate patients with high levels of expression; Fig. S8c) confirmed a significantly better OS (5-years OS 44.0% vs. 0%, p=0.017 in TCGA-LAML; and 39.8% vs. 20.8%, p=0.034 in Bohlander AML) (Fig. 7h), confirming findings in the ‘training’ cohort (Fig. 7g).

Taken together, the data support a predictive working model in which inflammatory remodeling of LEPR+ stromal niche cells suppresses normal hematopoiesis (via downregulation of HSPC regulatory factors), but in which leukemia relapse-initiating cells may remain critically dependent on residual levels of stromal niche support (in particular SCF/KITLG-cKIT signaling) for their survival in the context of chemotherapeutic treatment (Fig. S9a). Strong inflammation-associated loss of stromal niches may take away this support, resulting in the impediment of leukemia (initiating) cells and reduced risk of relapse (Fig. S9b). Alternatively, but not mutually exclusive, direct engagement of leukemia-relapse-initiating cells by an activated immune system (driving inflammatory alterations in stromal cells) may explain the association between inflammation and reduced relapse risk in these patients. Future experiments are warranted to test these working models based on our data.

## DISCUSSION

Tumor-promoting inflammation is considered an enabling characteristic of tumorigenesis via tumor-promoting effects that immune cells have on neoplastic cells and disease progression [1]. The exact mechanisms linking inflammation to oncogenesis, however, remain incompletely understood.

This, to our knowledge, is the first study to examine the predicted interactions between residual tissue resident stem/progenitor cells versus their neoplastic counterparts within their native niches and immune environment in *human* cancer at cellular resolution. By exploiting insights in the hematopoietic system, in which stem cells and their niches have been defined at near cellular resolution, we provide experimental support for the view that tumor-associated inflammation can result in the remodeling and deterioration of innate stem/progenitor cell niches. This is predicted to result in the loss of their capacity to support residual normal HSPCs with relative resistance of neoplastic cells, thus providing a conceptual basis for tissue repression and competitive advantage of neoplastic cells in AML (Fig. S9a).

The data provides human disease relevance to concepts previously postulated by experiments in various murine models, in which leukemic cells alter stromal niches in ways that were proposed to inhibit normal hematopoiesis (by suppression of key HSPC factors) [24] or secretion of inhibitory factors [22, 23], but at the same time remain dependent on these niches for their survival under chemotherapeutic conditions [16, 64]. Our human data implicate inflammatory signaling as a key driver of niche deterioration in AML with potential therapeutic significance.

It is important, however, to note that the cellular taxonomies we established do not include all niche cells that constitute the mammalian hematopoietic system. Specifically, endothelial cells and stromal niche subsets with close anatomical relationship to the (trabecular) bone, as identified in the murine bone marrow [24, 65, 66], are likely absent or underrepresented in human BM aspirates. The limited availability of core biopsies (not a universal diagnostic procedure in AML), as well as the low amount of available tissue and heterogeneity (sampling bias) of core biopsies preclude robust analyses of these types of niche cells presently.

The stromal cells we retrieved from human aspirates show transcriptional resemblance to the LEPR+ perivascular stromal HSPC niches identified in mice [5, 67]. This subset of stromal cells, thought to largely overlap with so called CXCL12-abundant reticular (CAR) cells, is considered to be the largest subset of stromal cells in the mammalian bone marrow [2] and have previously been demonstrated to be instrumental for the maintenance of normal HSCs and committed progenitors [68].

In addition to inflammatory remodeling of LEPR+ stromal niches, we describe transcriptional heterogeneity in the human HSC pool and report a previously unanticipated loss of a specific subset of HSCs/MPPs in AML that show strong transcriptional congruence with ST, high output HSCs previously defined in mice, with relative conservation of LT, low-output HSCs. These findings may provide a cellular basis for the longstanding observation that normal hematopoiesis is typically reconstituted after chemotherapeutic eradication of leukemic cells, which indicated that a population of cells with HSC characteristics must be able to survive these conditions. Our data indicate that LT-HSCs in the human bone marrow are relatively resistant to inflammatory stress and the loss of SCF from stromal niches, perhaps through the expression of inflammation inhibiting pathways such as the *NRA* family and *TNFAIP3* [36–38]. Other mechanisms likely play a role, and it may be that the transition from LT-HSCs to ST-HSCs is specifically impaired in human AML by yet to be identified factors in the leukemic bone marrow. The identification of transcriptional human HSC subsets and elucidation of the cellular taxonomy of AML reported here is anticipated to enable and facilitate future investigations in these directions.

The nature of the drivers of stromal inflammation remains largely speculative at this point. Our data demonstrate that TNFα, which is commonly overexpressed in myeloid malignancies, is able to induce inflammatory activation and remodeling of the stromal microenvironment. The AML taxonomy indicated that activated immune cells may be a source of TNFα in AML, reminiscent of recent work in mice demonstrating that, in viral infection, activated BM resident CD8^+^ T cells induce damage to stromal niches associated with activation of inflammatory pathways and reduction of expression of Cxcl12 and Scf [69]. It is, however, reasonable to assume that other sources of TNFα may exist and that multiple inflammatory cytokines are overexpressed in the AML environment that may contribute to stromal remodeling. Future investigations, instructed by the current taxonomies may shed further light on this. It is further important to note that the negative effects of TNFα on residual normal hematopoiesis in AML are unlikely to be entirely stromal niche-dependent, as TNFα has been demonstrated to exert both positive and negative direct effects on HSPCs [56, 70–73].

Inflammatory remodeling of stromal niches and the associated loss of niche support was associated with a favorable outcome upon chemotherapeutic treatment, supporting the notion that it may pose a therapeutic vulnerability for leukemia relapse-initiating cells that may remain critically dependent on some residual niche support to ascertain their survival during chemotherapeutic treatment (Fig. S9b). Alternatively, or in addition, inflammatory responses driven by activated innate and adaptive immune cells may contribute to the eradication of leukemia-relapse initiating cells following chemotherapy in niche-independent fashions.

The finding that niche signatures associated with clinical outcome in AML patients treated with intensive chemotherapy may have impact on risk stratification and therapeutic decision making in AML, in which current prediction models are instructed by parameters from hematopoietic cells, rather than their stromal environment [59].

Finally, the presented data is anticipated to provide an important resource of human bone marrow signaling, and in particular stem-cell niche interactions, defining normal and leukemic hematopoiesis, to serve as a platform for discovery and validation of findings from non-human model systems.

## METHODS

### Human normal and AML bone marrow samples

Normal bone marrow samples were obtained by hip bone aspiration from healthy donors for allogeneic transplantation.

AML bone marrow aspirates were obtained from AML patients (age 18-65) included in the HOVON-132 clinical trial testing the addition of lenalidomide to intensive treatment in younger and middle-aged adults with newly diagnosed AML [57]. Patients received two cycles of intensive induction chemotherapy. Cycle 1 included idarubicin at 12 mg/m^2^ (3-hour infusion on days 1, 2, and 3) and cytarabine at a dose of 200 mg/m^2^ (per continuous infusion on days 1-7) with or without lenalidomide. Cycle 2 contained daunorubicin 60 mg/m^2^ per 1-hour infusion on days 1, 3, and 5 plus cytarabine 1000 mg/m^2^ given intravenously for 3 hours twice per day on days 1 to 6 with or without the addition of lenalidomide. Patients in CR or CR with incomplete hematologic recovery (CRi) after cycle 2 received consolidation with 1 final additional cycle of intensive chemotherapy with mitoxantrone-etoposide (cycle 3), autologous stem cell transplantation (auto-SCT) or allogeneic stem cell transplantation (allo-SCT). The study was approved by the ethics committees of the participating institutions and was conducted in accordance with the Declaration of Helsinki. All patients gave their written informed consent [57]. Bone marrow specimens were collected by hip bone aspiration at diagnosis, after remission cycle 2 and 3-6 months after auto/allo-SCT.

Mononuclear cell fractions of human bone marrow aspirates were isolated using lymphoprep, and viably frozen in PBS supplemented with 40% heat-inactivated fetal calf serum (FCS; Corning) and 10% dimethyl sulfoxide (DMSO; Sigma-Aldrich). Marrow aspirates for scRNAseq were cryopreserved within 24 h after collection. All specimens were collected with informed consent, in accordance with the Declaration of Helsinki.

### Patient cell isolation for RNA sequencing

Viably frozen bone marrow aspirates (mononuclear cell fractions) were thawed in a water bath at 37°C and washed with warm Dulbecco’s Modified Eagle Medium (DMEM) supplemented with 10% fetal calf serum (FCS) as described in the 10X Genomic protocol ‘Fresh Frozen Human Peripheral Blood’.

For isolation of hematopoietic fractions, 10% of the thawed bone marrow cells (>20 x 10^6^) from each individual were stained for sorting in PBS supplemented with 0.5% FCS at 4 °C with the following antibodies: CD45-APC (1:20, clone 2D1; eBioscience), CD34-AF700 (1:50, clone 581; Biolegend), CD117-PE-CF594 (1:50, clone YB5.B8; BD Biosciences), CD33-PE (1:50, clone P67.6; BD Bioscience), CD3-PE-Cy7 (1:50, clone SK7; Biolegend), CD19-APC-Cy7 (1:50, clone HIB19; Biolegend), CD38-FITC (1:50, clone HIT2; Life Technologies). For exclusion of dead cells, 7AAD (1:100; Beckman Coulter) was used. The 7AAD^-^CD45^+^CD34^+^ HSPC fraction, 7AAD^-^CD45^+^CD34^-^CD117^+^ and 7AAD^-^CD45^+^CD34^-^CD33^+^ myeloid fraction, and 7AAD^-^CD45^+^CD34^-^CD117^-^CD33^-^ non-myeloid (lymphoid) fraction were sorted in DMEM supplemented with 10% FCS using a FACSAria III (BD Biosciences) and BD FACSDiva version 5.0.

The non-hematopoietic fraction was sorted from the thawed bone marrow cells as described before [74]. 90% of the thawed cells (>200 x 10^6^) were stained with biotinylated antibodies against CD45 (1:50, clone HI30; Biolegend) and CD235a (1:50, clone HIR2, Biolegend) followed by depletion using magnetic anti-biotin beads (20 µl per 10^7^ cells; Miltenyi Biotec) in PBS supplemented with 2% FCS and iMag (BD Biosciences). After depletion, the remaining cells were stained for sorting in PBS containing 0.5% FCS at 4 °C with the following antibodies: Streptavidin-AF488 (1:100, Invitrogen), CD45-BV510(1:50, clone HI30; Biolegend), CD235a-PE-Cy7(1:50, clone HI264; Biolegend), CD71-AF700 (1:20, clone MEM-75; Exbio), CD271-PE (1:50, clone ME20.4; Biolegend), CD105-APC (1:50, clone SN6; eBiosciences), CD31-APC-Cy7 (1:20, clone WM59; Biolegend), CD34-eFluor610 (1:50, cone 4H11; eBioscience) and CD144-V450 (1:50, clone 55-7H1; BD Biosciences). 7AAD (1:100; Beckman Coulter) was used for dead cell exclusion. FACSAria III (BD Biosciences) and BD FACSDiva version 5.0 were applied for sorting. 7AAD^-^CD45^-^CD235A^-^CD71^-^ non-hematopoietic fraction was sorted in DMEM supplemented with 10% FCS for Single-cell RNA sequencing and 7AAD^-^CD45^-^CD235A^-^CD71^-^CD31^-^CD274^+^ BMSCs were sorted in TRIzol (Life Technologies) for bulk RNA sequencing.

### Single-cell RNA sequencing

Prior to single-cell RNA sequencing, two fractions sorted separately from the same donors were pooled together (non-hematopoietic fraction with lymphoid fraction and myeloid fraction with HSPC fraction) followed by encapsulation for barcoding and cDNA synthesis using the Chromium Single-cell 3′ Reagent kit v3 (10x Genomics). 3’ gene expression library was constructed according to the manufacturer’s recommendations. The quality and quantity of libraries was determined using an Agilent 2100 Bioanalyzer with 2100 Expert version B.02.11.SI811 software and a High Sensitivity DNA kit. Libraries were sequenced on a NovaSeq 6000 platform (Illumina), paired-end mode, at a sequencing depth around 45,000 reads per cell, followed by computational alignment using CellRanger (version 3.0.2, 10x Genomics).

Seurat (R package, version 4.0.0) [75] was used for data preprocessing and downstream analysis. For data preprocessing, datasets were first subjected to quality control steps that included removing doublets (a high ratio of RNA counts vs. feature numbers (> 6)) and filtering out apoptotic cells determined by high transcriptional output of mitochondrial genes (>5% of total). Subsequent, single cell data from separate runs of the same donor were merged to generate a complete picture that includes all cell types for each individual. Aiming at generating a tissue map robustly representing the bone marrow taxonomy in health individuals and AML patients, the merged datasets of each individual were integrated using the integration function in Seurat followed by linear dimensional reduction. That included scaling of gene expression across all cells, principle component analysis (PCA) on the most variable genes (*k*L=L2,000) and unsupervised clustering using a shared nearest neighbor (SNN) modularity optimization-based clustering algorithm (resolution 0.3–1).). To mitigate the effects of cell cycle heterogeneity, cell cycle phase scores were calculated based on canonical markers, and were regressed as described in Seurat protocol. The data was visualized using Uniform Manifold Approximation and Projection for Dimension Reduction (UMAP) [76] (McInnes et al., 2018. Cell types were identified by the Clustifyr R package [25] and by reviewing expression of canonical markers associated with particular cell types. In order to reveal BMSC heterogeneity at a high resolution, BMSCs were separated using the Cellselector function in Seurat and independently pre-processed/analyzed. For HSC analysis, the HSC/MPP population was re-clustered in order to study HSC heterogeneity robustly in single samples. Only the samples with >10 cells in the HSC/MPP population were used for the analysis.

Cell trajectory analysis was performed using Monocle3 R package (https://github.com/cole-trapnell-lab/monocle3). Cellchat and CellphoneDB were applied to predict ligand-receptor interaction between cell types [39, 77]. For differential expression gene analysis between samples, a DEseq2 [78] based *pseuD*E package was developed for the aggregation of single-cell level gene-counts and application of differential expression (DE) analysis (software is available at https://github.com/weversMJW/pseude). For Gene set enrichment analysis (GSEA) [79] between groups, variable genes were identified using Seurat‘s Findmarker function and ranked using the formula *–sign(log2foldchange)* log10(p-value)*. The R package fgsea was used for the analysis with a permutation of 1000 using predefined gene sets from the Molecular Signatures Database (MSigDB 6.2) as input. Gene enrichment scores for individual cells were calculated using Seurat’s AddGeneScore function which calculates the score by counting the average expression levels of provided genes, subtracted by the aggregated expression of randomly selected control genes.

### Identification of *NPM1m* cells

The *NPM1* mutation is a 4-nucleotide insert located in the last exon proximal to its 3’- end. The enzyme encoded by *NPM1* plays an important role in ribosome biogenesis and functions as protein chaperone. Hence, its expression is ubiquitous across all captured cell types and is specifically high in cells of hematopoietic origin. The high expression of *NPM1* and the position of the driver mutation towards the 3’-terminus makes this mutation a prime candidate for detection in 3’ single cell transcriptional data.

Identification of cells carrying the *NPM1* mutation was performed using an in-house developed tool. In short, alignment results produced by the Cell Ranger pipeline in the form of a BAM file was used as input. Aligned reads, representing sequenced cDNA molecules, were extracted at the mutation position and screened for the *NPM1* mutation taking into account the cellular barcode (CB) and unique molecular identifier (UMI), irrespective of whether the cell is of hematopoietic origin. Following UMI-based consensus sequence building, mutant-carrying reads were assigned to the cell of origin based on the CB. Reads without direct detection of the *NPM1* mutation, i.e. the known 4-nucleotide insert is absent, were screened by a second approach. The *NPM1* mutation is located close to an intron-exon boundary which can result in improper alignment when the mutation is positioned near the extremities of the cDNA molecule, i.e. the 4-nucleotide insert mutation is partially, abnormally or not introduced (only missense variants) in the aligned read, which in turn complicates detection of the mutation. To prevent this issue all reads without detected *NPM1* mutation in the first screening round were aligned to the reference genome (hg38) and a modified version thereof including the 4-nucleotide insert mutation, in both cases taking into account the local *NPM1* splicing pattern (i.e., splicing from the penultimate to terminal exon). Reads aligning best to the reference genome were labeled wildtype and those aligning better to the modified version were labeled mutant. Using UMI-based consensus building, mutant-carrying and wildtype reads identified in the second round of screening were attributed to cells based on the CB. Subsequent to the two screening round each cell has assigned a number of UMI-controlled wildtype and mutant reads covering the mutation position. Cells with at least one mutant read detected were classified as *NPM1* mutant. Cells with at least 3 wildtype reads detected covering the mutation position are classified as likely *NPM1* wildtype. All remaining cells that did not pass these criteria were classified as ‘non assignable (NA)’. Software is available at: https://github.com/RemcoHoogenboezem/annotate_bam_statistics_sc.

### Bulk RNA sequencing

RNA was extracted using TRIzol reagent (Invitrogen) according to the manufacturer’s instructions, in combination of isolation with GenElute LPA (Sigma-Aldrich). cDNA was prepared using the SMARTer procedure through the SMART-Seq v4 Ultra Low Input RNA Kit (Clontech) for Illumina Sequencing. Quantity and quality of cDNA production was assessed using the Agilent 2100 Bio-analyzer and the High Sensitivity DNA kit. cDNA libraries were generated using the TruSeq Sample Preparation v2 guide (Illumina) and paired-end sequencing on a NovaSeq 6000 (Illumina). Adaptor sequences and polyT tails were trimmed from unprocessed reads using fqtrim version 0.9.7. (http://ccb.jhu.edu/software/fqtrim/). Read counts and TPM (Transcripts Per Million) were determined with Salmon version 1.2.1 [80]. Gene count estimates were determined from Salmon output with the tximport R package version 1.16 [81]. These gene count estimates were in turn normalized, prefiltered according to standard practices and used for determining the differential gene expression between groups of interest through the DESeq2 package (version 1.28) [78] using default parameters. GSEA was performed with GSEA software (version 3.0, Broad Institute) using predefined gene sets from the Molecular Signatures Database (MSigDB 6.2). Gene lists were ranked on the basis of the log2LFC made available through the DESeq2 package. Classical enrichment statistics with 1,000 permutations was used to determine significant enrichment within gene sets. GSVA (R package) was applied for gene enrichment analysis in single samples.

### Fluorescence Immunohistochemistry

Bone marrow biopsies were fixed in 4% paraformaldehyde, decalcified in 150 mM EDTA, washed in 70% ethanol and embedded in paraffin. Sections of 5-μm bone marrow biopsies were deparaffinized in xyleen and hydrated in a graded series of ethanol. Antigen retrieval was achieved by microwave treatment in TRIS-EDTA buffer (1M Tris, 0.5M EDTA, 0.05% Tween-20, PH=9.0). Slides were subsequently blocked using BloxALL (Vector laboratories), to block endogenous peroxidases, PBS supplemented with 0.5% Tween-20, 5% human serum, 5% goat serum and 5% donkey serum to circumvent aspecific binding, and were stained overnight at 4°C with primary antibodies. The primary antibodies that were used in this study included rabbit anti-human CD271 (1:100, HPA004765, Biolegend), mice anti-human CXCL12 (1:50, clone 79018, Invitrogen), and rat anti-human CD44 (1:50, clone IM7, Invitrogen). The secondary antibodies included alexa fluor 488 labelled donkey anti-mouse (xxx, 1:200), horseradish peroxidase (HRP)-labeled goat anti–rat (1:100, Jackson ImmunoResearch), and cy5 labeled donkey anti-rabbit antibodies (1:800, Jackson ImmunoResearch).. An Tyramide superboost kit (Invitrogen) was used to label CD44 in AF555 following the manufacturer’s protocol. The stained sections were mounted in ProLong Diamond containing DAPI (Invitrogen). Images were acquired using a Leica SP5 confocal microscope at 40X magnification and were subsequently analyzed using Fiji software.

### Animal experimental procedures

3 month old C57BL/6 mice were purchased from Charles River and maintained in specific pathogen free conditions in the Experimental Animal Center of the Erasmus MC (EDC). These mice were intraperitoneally (I.P.) injected with 5µg recombinant murine TNFα (Peprotech) for 5 consecutive days, followed by peripheral blood collection for complete blood count, and were sacrificed at day 6 by cervical dislocation. For establishment of *MLL-AF9* AML mice model, retroviral plasmid containing MLL-AF9- EGFP (a gift from Dr. Stefan Erkeland) were transfected into the platinum-E (Plat-E) retroviral packaging cell line using LipoD293 transfection reagent (SignaGen) for retroviral production. The virus was transduced into freshly isolated Lin^-^ bone marrow cells derived from 3 month old B6.SJL mice (from Charles River) after Lin^+^ cell depletion using mouse Lineage Cell Depletion Kit (Miltenty Biotec). 1 million transduced cells (containing ±5% GFP^+^ cells) were intravenously injected into lethally irradiated (9.5Gy) age-matched C57BL/6 mice. When AML had developed (white blood cell counts >15*10^3^/mm^3^) 30 days after injection, mice were sacrificed and bone marrow cells (%GFP^+^>90%) were collected. For testing effects of TNFα on AML, 50,000 bone marrow cells derived from the MLL-AF9 AML mice were intravenously injected into non-irradiated 3 month old C57BL/L mice. After 30 days of transplantation, 5ug recombinant murine TNFα (Peprotech) was I.P. injected into the mice for 5 consecutive days, followed by peripheral blood collection for complete blood count. Mice were sacrificed at day 11 (counted from the start date of TNFα injection) by cervical dislocation. Animal studies were approved by the Animal Welfare/Ethics Committee of the EDC in accordance with legislation in the Netherlands (approval No. EMC 2067, 2714, 2892, 3062).

### Flow cytometry for mice bone fractions and HSPCs

Mice collagenased bone fraction cells were stained with CD45.2-APC-Cy7 (1:200, clone 104, eBiolegend), Ter119-BV510 (1:50, clone TER-119, BioLegend), CD31-PE-Texas- red (1:100, clone MEC13.3, BD Bioscience), CD144-PE-Cy7 (1:200, clone BV13, Biolegend), CD51-PE (1:50, RMV-7, Biolegend), Sca1-PacificBlue (1:100, D7, Biolegend) and LEPR-biotin (1:50, catalog RB01, R&D systems) followed by staining of streptavidin-APC (1:100, Biolegend) and 7AAD (1:100) in PBS + 0.5% FCS. 7AAD^-^CD45.2^-^Ter119^-^CD31^-^CD114^-^CD51^+^Sac1^-^LEPR^+^ and 7AAD^-^ CD45.2^-^Ter119^-^CD31^-^CD114^-^CD51^+^Sca1^-^LEPR^-^ bone marrow stromal cells were sorted in TRIzol using FACSAria III (BD Biosciences) and BD FACSDiva version 5.0. Data was analyzed using FlowJo software. For HSPC analysis, mice bone marrow mononuclear cells were first stained with lineage (Lin) cocktail containing biotinylated antibodies against Ly-6G/Ly-6C (Gr-1) Biotin RB6-8C5 (BD Bioscience) (1:100), CD11b Biotin M1/70 (BD Biosciences) (1:100), Ter119 Biotin TER-119 (BD Biosciences) (1:100), CD3e Biotin 145-2C11 (BD Biosciences) (1:100), CD4 Biotin GK1.5 (BD Biosciences) (1:100), CD8 Biotin 53-6.7 (BD Biosciences) (1:100) and B220 Biotin RA3-6B2 (BD Biosciences) (1:100). After one washing step, cells were incubated with Streptavidin Pacific Orange (Life Technologies) (1:200), together with a combination of the following antibodies: Sca1 Pacific Blue E13-161.7 (BioLegend) (1:100), CD48 AF700 HM48-1 (BioLegend) (1:100), CD150 PE-Cy7 TC15-12F12.2 (BioLegend) (1:100), CD127(IL7RA) APC A7R34 (Biolegend) (1:100), c-Kit PE-CF594 2B8 (BD Biosciences) (1:100), CD16/32 APC-Cy7 2.4G2 (BD Biosciences) (1:100), FLT3 APC A2F10 (1:40) and 7-ADD (1:100).

### Cell line and ex vivo co-culture

The human BMSC cell line HS-5 was purchased from ATCC and maintained in RPMI- 1640 medium supplemented with 10% FCS and 1% Pen-strep. Primary CD34^+^ cells were obtained from umbilical cord blood (CB) using a Ficoll gradient protocol and by magnetic-activated cell sorting (MACS). Primary AML cells were obtained from patients’ aspirates as described above. For ex vivo co-culture, 180.000 HS-5 cells were seeded in each well of a 12-well plate in RPMI-1640 medium supplemented with or without 10ng/ml recombinant human TNFα. After 24 hours and washing of HS-5 cells with PBS, CB CD34^+^ cells or primary AML cells were seeded on top of the HS-5 cells and covered with GMP serum-free Stem Cell Growth Medium (CellGenix GmbH) supplemented with 50ng/ml TPO, 50ng/ml FLT3, 50ng/ml SCF, and 20 ng/mL IL3 (only for AML cells). Cells were cultured for 3 days at 37°C with 5% CO2. All cytokines used in the experiment were purchased from Peprotech.

Counting of CB CD34^+^ cells after 3 days of ex vivo co-culture was accurately obtained with flow-count fluorosphere beads (Beckman Coulter), in combination with the following antibodies: Lin-cocktail-FITC (1:100, catalog 22-7778-72, eBioscience), CD45-APC (1:20, clone 2D1; eBioscience), CD34-AF700 (1:50, clone 581; Biolegend). For counting of AML cells, an antibody cocktail containing CD45-APC (1:20, clone 2D1; eBioscience), CD34-AF700 (1:50, clone 581; Biolegend), CD117-PE-CF594 (1:50, clone YB5.B8; BD Biosciences), CD33-PE (1:50, clone P67.6; BD Bioscience) was used in combination with flow-count fluorosphere beads. In both cases, dead cells were excluded based on the DAPI (1:7500) gate. The data was acquired using a LSRII flow cytometer (BD Biosciences) and analyzed using FlowJo software.

### CFU-GEMM assay

On day 3 of co-culture, 2000 mononuclear cells (MNCs) from HS-5-Veh (PBS) condition and in HS-5-TNFα condition were resuspended in IMDM medium. This cell suspension was mixed with MethoCult™ GF H84434 (StemCell Technologies), which allows the growth of colonies from all three lineages, and triplicate dishes were plated. The methocult plates were kept at 37°C in a 5% CO2 incubator for 2 weeks until colony counting under a light microscope (Zeiss).

### Survival analysis

The R package dCVnet (https://github.com/AndrewLawrence/dCVnet) was used to perform Lasso-penalized nested cross-validated Cox regression (k-fold.outer = 5, k-fold.inner = leave-one-out cross-validation [i.e., 1]) to build a predictive model for overall survival (OS) based on the centered and scaled gene expression of the 189 genes from all available AML samples (n=62). This resulted in the retention of 13 out of 189 genes weighted according to their contribution to the model. Similar to Ng. et al [61], we obtained a weighted score by taking the linear combination of the centered and scaled gene expression levels of the 13 retained genes weighted by the obtained regression coefficients. Score = (*ADAMTS4* x -0.066) + (*ZBTB21* x -0.079) + (*HAS2* x -0.035) + (*SPAG9* x -0.104) + (*KITLG* x 0.345) + (*CXCL3* x 0.021) + (*MEDAG* x 0.224) + (*PTGDS* x 0.094) + (*ARPC5L* x 0.269) + (*UHRF1BP1L* x -0.0008) + (*SMAD7* x -0.069) + (ANXA2 x 0032) + (ELL2 x -0.072). The median score value was used to dichotomize the AML cohort into low and high score groups. These groups were labelled as ‘BMSC niche preserved’ (low score) and ‘BMSC niche inflammatory remodeling’ (high score), respectively. Distinction into the ‘favorable’, ‘intermediate’ and ‘adverse’ groups was based on the ELN2017 genetic risk classification. The log-rank test was used to assess statistical differences between the survival distributions, a p-value ≤ 0.05 was considered statistically significant. For clinical outcome analysis using public-available datasets, TCGA-LAML and Bohlander AML (GSE37642) datasets were acquired using R package TCGAbiolinks and GEOquery, respectively. Patients between 18-65 years old were selected and stratified into KITLG low and KITLG high group based on 75% percentile of normalized counts of KITLG. The log-rank test was used to assess statistical differences between the survival distributions, a p-value ≤ 0.05 was considered statistically significant.

### Statistics

Statistical analysis was performed using Prism 8 (GraphPad Software) and/or R program. Unless otherwise specified, unpaired, two-tailed Student’s t test (single test), one-way ANOVA (multiple comparisons) or Spearman’s rho test (correlation analysis) were used to evaluate statistical significance, defined as p-value (pval) < 0.05. All results in bar graphs are mean value ± SD.

### Author contributions

Conceptualization: M.H.G.P.R.; Methodology: L.C, E.P, C.v.D, J.F, T.v.T, M.Y, P.P, M.J, A.B, R.H, C.W, E.B, B.L, T.C, M.A.S. and M.H.G.P.R; Investigation: L.C, E.P, C.v.D, J.F, T.v.T, M.Y, P.P, M.J, A.B, R.H, C.W, E.B, M.A.S; Resources: B.L, M.J. T.C; Data Curation: M.A.S., R.M.H.; Writing: L.C, M.A.S. and M.H.G.P.R.; Visualization: L.C, E.P, C.v.D, M.A.S, M.H.G.P.R.; Supervision and Funding Acquisition: M.H.G.P.R.

## Acknowledgements

The authors wish to thank Nathalie Papazian and Michael Vermeulen for technical assistance, Mariette ter Borg for providing CD34+ cells; the HOVON/SAKK leukemia working group and all its members and participating sites for conducting the HOVON132 trial. Peter Valk, Patrycja Gradowska and Jurjen Versluis for providing the genetic and clinical data. The Josephine Nefkens Precision Cancer Treatment Program for infrastructural support and members of the Erasmus MC Department of Hematology for providing scientific discussion and members of the Erasmus MC animal core facility EDC for help with animal care.

## SUPPLEMENTARY FIGURE LEGENDS

**Figure S1.**
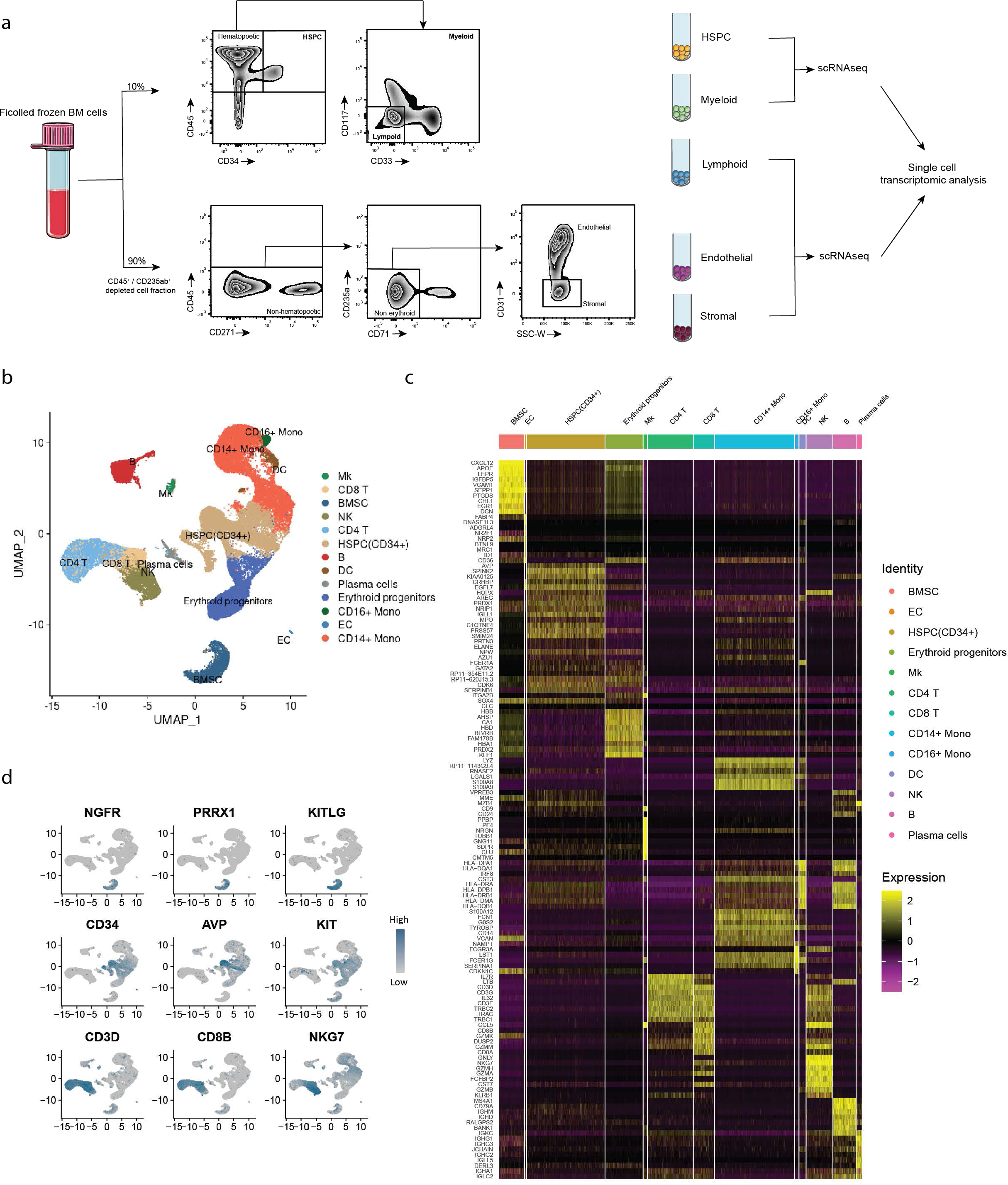
Experimental design and transcriptional identification of distinct cell types in the human normal bone marrow. (a-b) FACS gating strategy for sorting of hematopoietic and non-hematopoietic cell populations from bone marrow aspirates for scRNAsequencing. Bone marrow cells were subsorted into HSPC (CD45^+^CD34^+^), non-lymphoid/myeloid (CD45^+^CD34^-^ CD117^+^CD33^+^), lymphoid (CD45^+^CD34^-^CD117^-^CD33^-^), endothelial/non-stromal (CD45^-^ CD235a^-^CD71^-^CD31^+^) and stromal niche (CD45^-^CD235a^-^CD71^-^CD31^-^) fractions to obtain robust representation of all bone marrow cell types in the scRNAseq data. Sorted cells were pooled into two populations for scRNAseq, followed by integration of the data into a single ‘tissue map’ of the bone marrow. Clustifyr is used for Identification of cell types based on gene expression (Fu et al, 2020) and result was plotted by UMAP (b). (c) Heatmap of the highest differentially expressed genes within each cell population. (d) Expression of selected genes in the BMSC, HSPC and T/NK cell gates, respectively.

**Figure S2.**
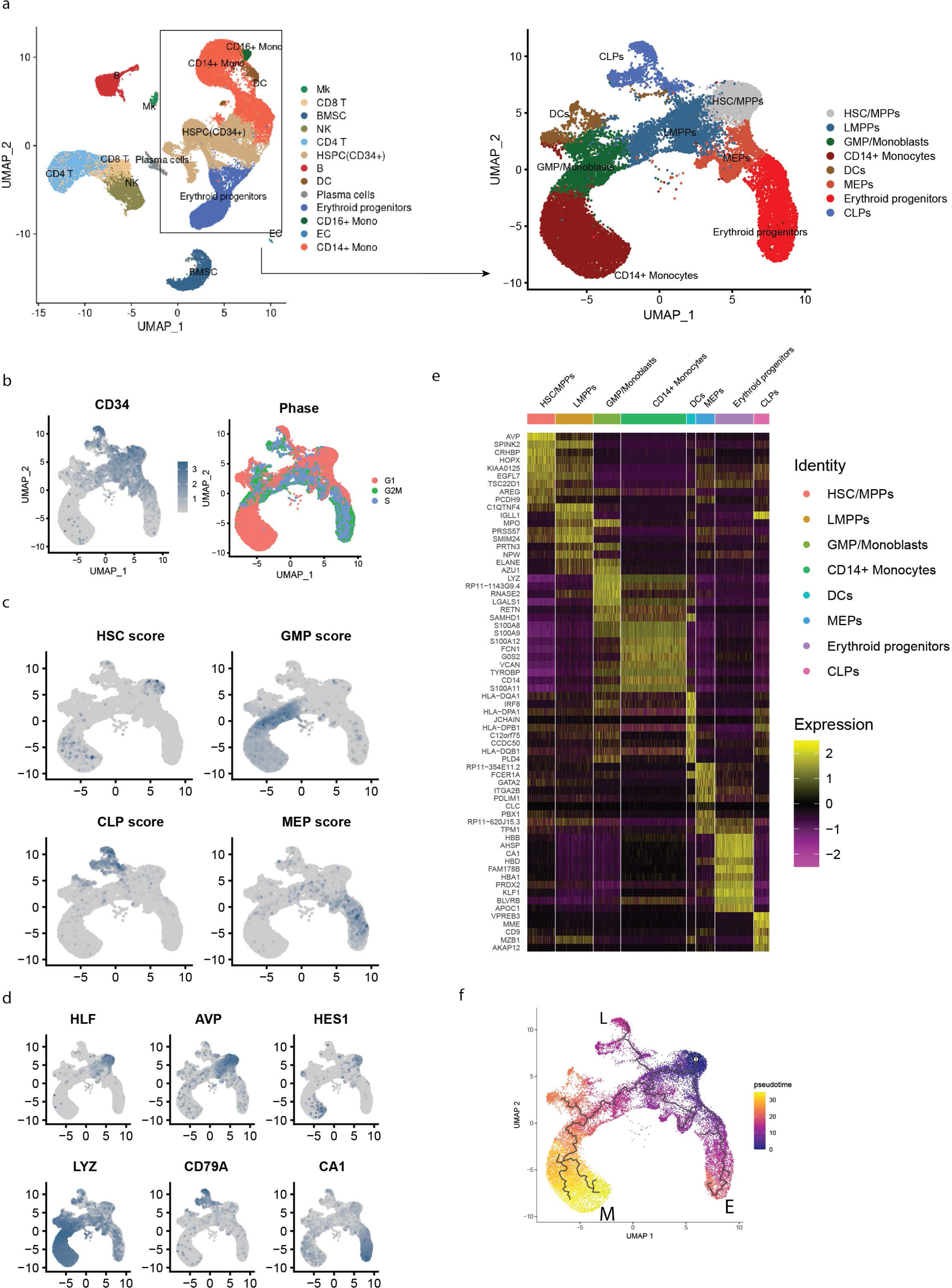
Transcriptional identification of HSPC subsets in the human normal bone marrow. (a-d) Re-clustering of the HSPC and myeloid population, and annotation into distinct subsets based on (b) CD34 expression and expression of cell cycle-defining gene sets, (c) gene score of HSPC types and (d) expression of genes indicative of HSPC types. (e) Heatmap of highest differentially expressed genes within each subpopulation. (f) Predicted lineage progression from HSCs to committed progenitor subsets by using pseudotime trajectory analysis. Calculated pseudotime is represented by color scale. L, lymphoid lineage. M, myeloid lineage. E, erythroid lineage.

**Figure S3.**
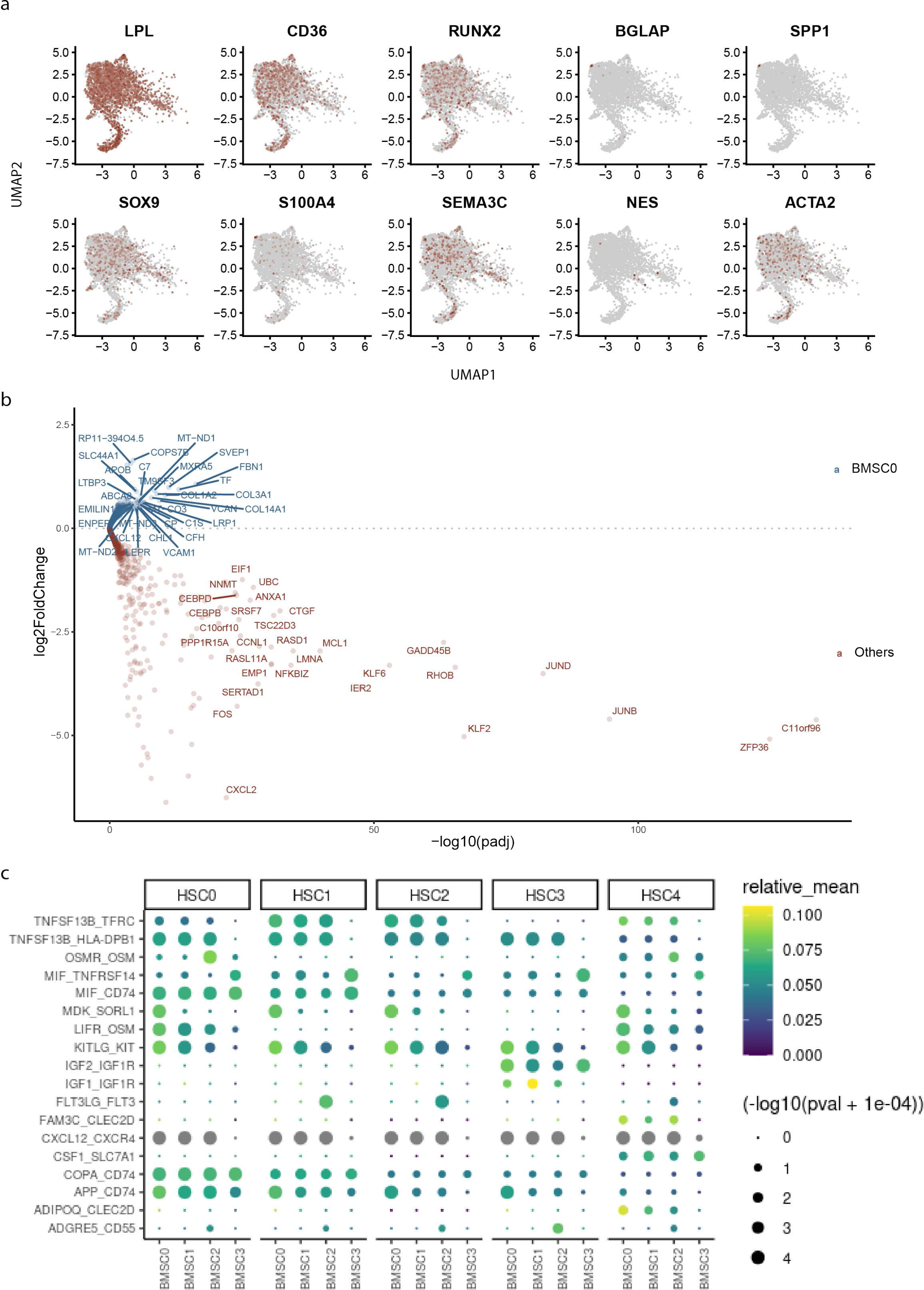
Transcriptional characterization of BMSC subsets in the human normal bone marrow. (a) Expression of genes related to adipocytic, osteolineage, chondrocyte, fibroblast and pericyte differentiation in BMSC subsets. (b) Volcano plot displaying genes differentially expressed in the BMSC-0 subset in comparison to other BMSC clusters (padj<0.05). Differential gene expression analysis is performed using the pseudoDE R package at sample level (pair-wise comparison in individual samples). (c) Predicted relative strength of ligand-receptor signaling pathways originating from BMSC subtypes to HSC/MPP subpopulations. Color scale and dot size represent relative mean strength and p-value of interactions, respectively. CellphoneDB was used for the assessment.

**Figure S4.**
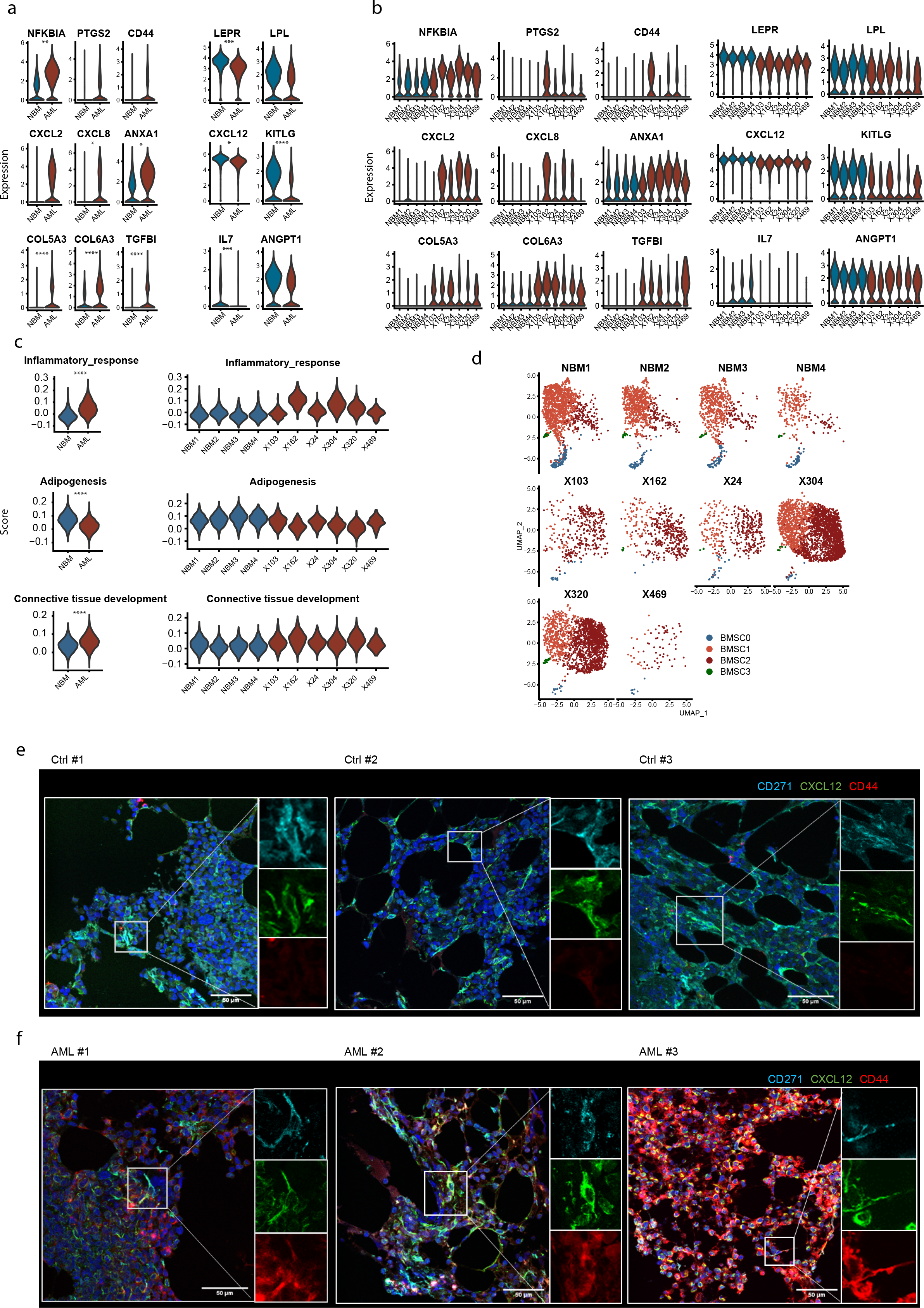
Remodeling of bone marrow stromal cells in *NPM1*m AML. Consistency within AML patient samples. (a) Comparison of expression of inflammation-associated genes, ECM remodeling-associated genes, pre-adipocytic markers and HSPC supportive factors in BMSC population. *, padj<0.05; **, padj<0.01; ***, padj<0.001. PseudoDE R package is used for differential expression gene analysis at sample levels. (b) Expression of inflammation-associated genes, ECM remodeling-associated genes, pre-adipocytic markers and HSPC supportive factors in BMSC population across all individuals. (c) Scores of Inflammation-, Adipogenesis-, and Connective Tissue Development-associated transcriptomic signatures in BMSC subsets across all individuals. (d) UMAP distribution of BMSC subsets across all individuals (4 healthy donors vs. 6 AML patients) demonstrating a consistent relative reduction of HSPC supportive cluster- 0 and a relative expansion of inflammatory cluster-2. (e-f) In situ inflammation of BMSCs in AML as demonstrated by expression of CD44 using fluorescence immunohistochemistry on bone marrow biopsies. Scale bar= 50µm.

**Figure S5.**
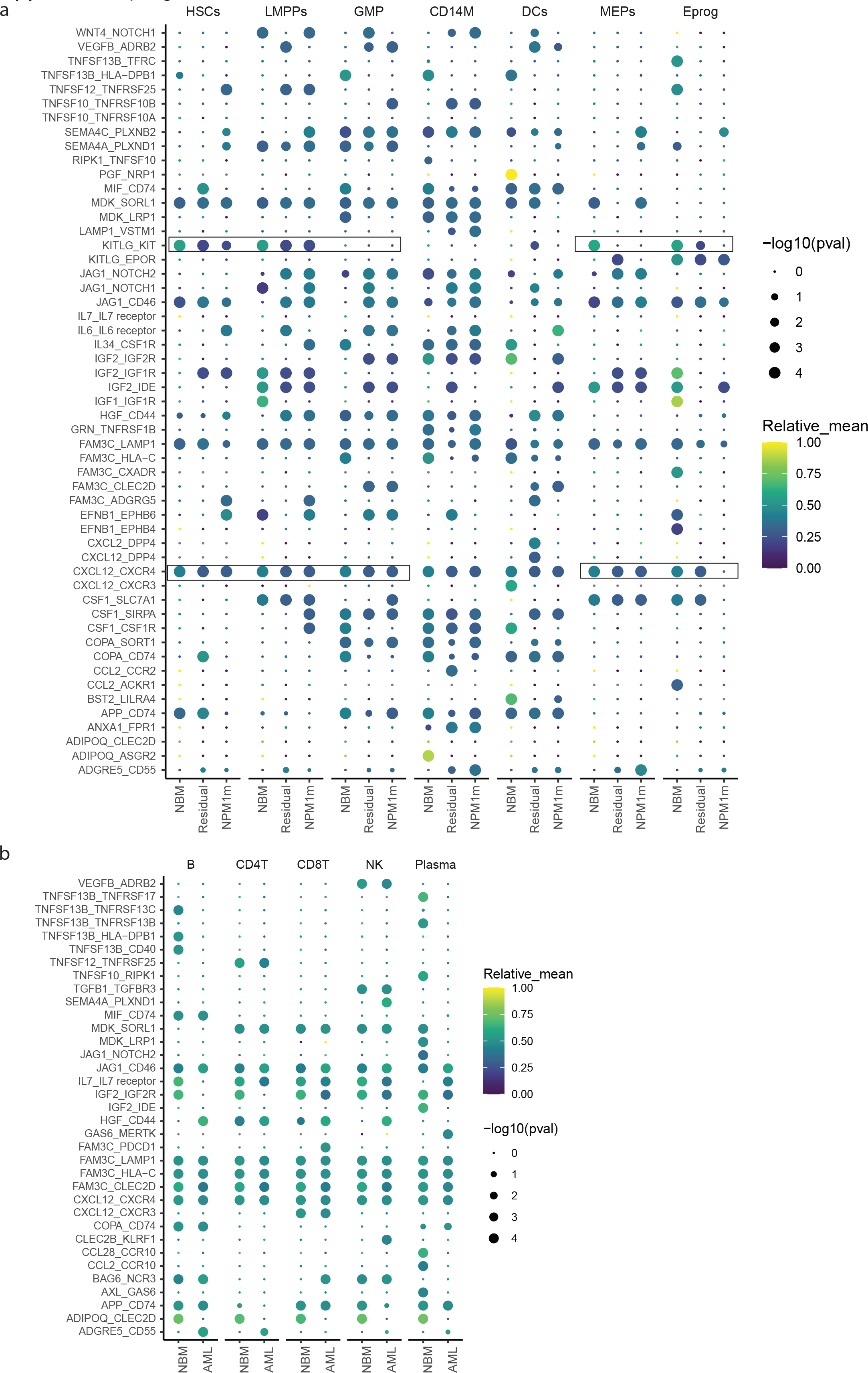
Predicted ligand-receptor interactions between BMSCs and other bone marrow resident cells in NBM and AML. (a) Predicted ligand-receptor interactions (CellphoneDB) from BMSCs to HSPCs in NBM, residual normal and *NPM1m* HSPCs. Note the reduced interactions of KITLG and CXCL12 with residual normal HSPCs in comparison to those in the NBM. Color-scale and size represent relative mean strength and p-value of the interactions. (b) Predicted ligand-receptor interactions from BMSCs to lymphoid immune cells, including CD4^+^ T cells, CD8^+^ T cells, NK cells, B cells, and plasma cells. Color-scale and size represent relative mean strength and p-value of the interactions.

**Figure S6.**
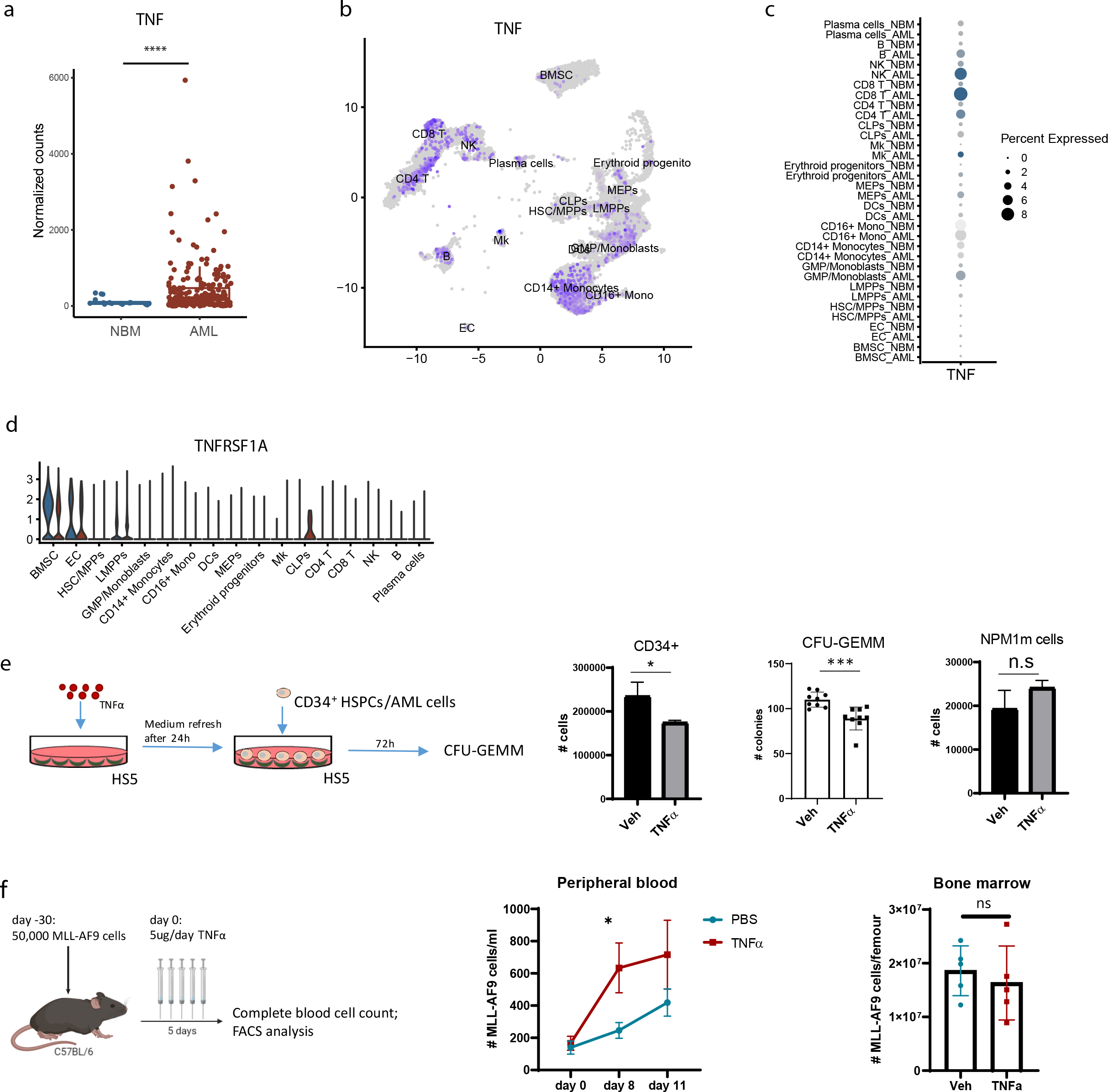
TNF**α** from adoptive and/or innate immune cells in AML is predicted to repress normal hematopoiesis via stromal niche signaling. (a) Overexpression of the gene encoding TNFα in AML in comparison to NBM. Data from the BEAT-AML cohort. n= 206 for AML and n=17 for NBM. ****, p<0.0001 (b-c) Relatively high expression of *TNF* in CD8^+^ T cells, NK cells and myeloid lineage cells in AML as shown in the UMAP plot (b) and dot plot (c). Size and color of the dot plot represent percentage and detectable expression level of TNFα. (d) Expression of the canonical TNFα receptor *TNFRSF1A* in bone marrow cell types in NBM and AML. Note that TNFRSF1A has the highest expression in the BMSC population. (e) Inflammatory activation of HS-5 human stromal cells by TNFα results in attenuation of normal hematopoiesis with relative resistance of *NPM1*m AML cells. HS-5 was pre-stimulated with 10ng/ml TNFα for 24 hours followed by co-culture with human cord blood (CB) CD34^+^ cells and *NPM1m* AML cells for three days (left). Numbers of CD34^+^ cells, CFU-GEMM (middle panel) and number of AML cells were counted. n=3. *, p<0.05. ***, p<0.001. CFU-GEMM: Colony forming unit for granulocyte, erythrocyte, monocyte and megakaryocyte. (f) Resistance of leukemic cells to TNFα exposure in a mouse model of AML. Administration of TNFα for five days did not result in reduction of leukemic cell number in the peripheral blood or bone marrow.

**Figure S7.**
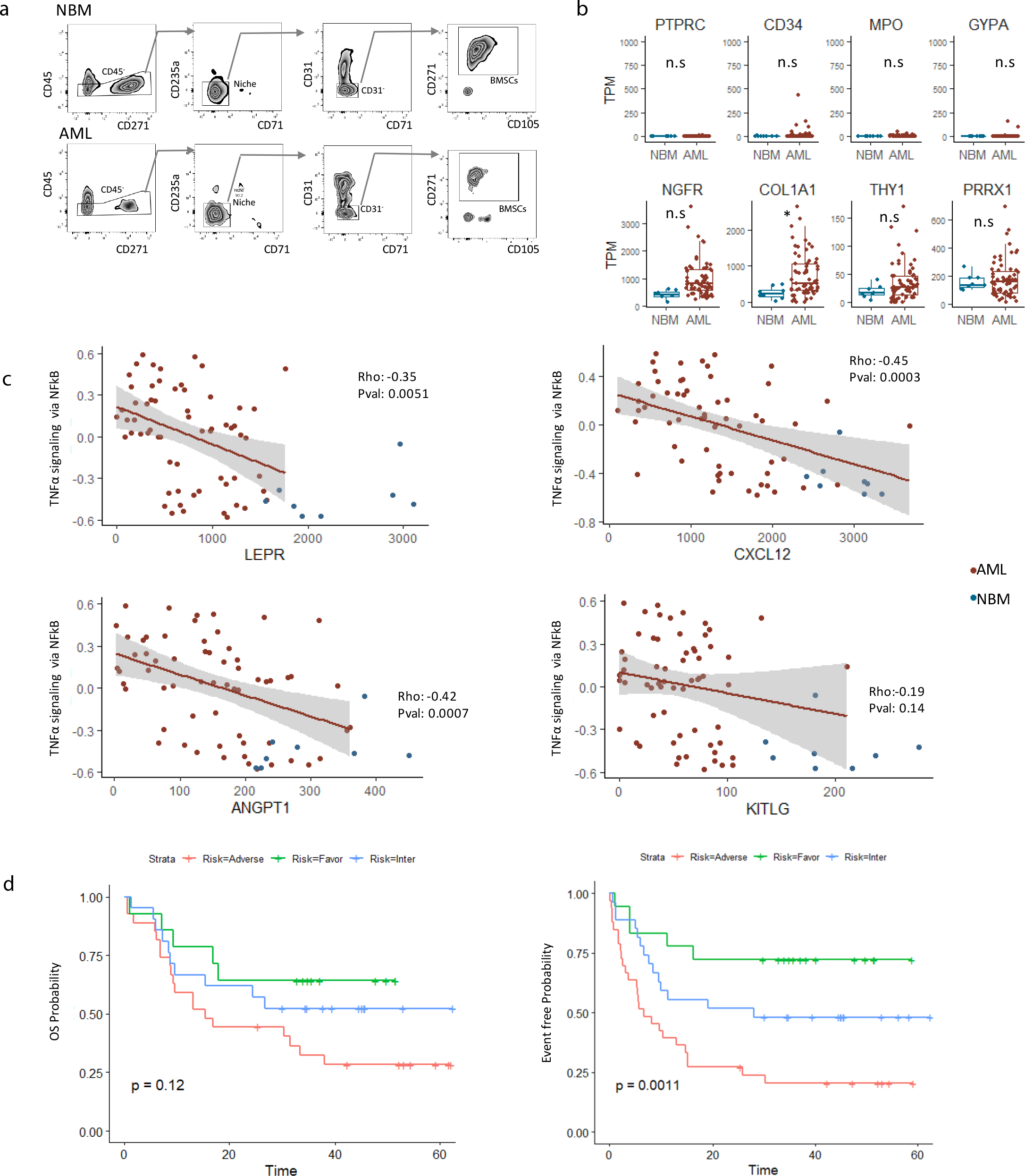
(NFkB-dependent) inflammatory activation and loss of HSPC factor expression in BMSCs are biologic commonalities in AML. (a) Representative FACS plots for sorting of CD45^-^CD235a^-^CD71^-^ CD31^-^ CD105^+^CD271^+^ BMSCs from NBM and AML for bulk RNA sequencing. (b) Expression of hematopoietic (upper panels) and stromal markers (lower panels) in BMSCs from NBM and AML. TPM = Transcripts per million. (c) Transcriptional activation of inflammatory programs in AML BMSCs is associated with reduced expression of niche/HSPC regulatory genes as demonstrated by Spearman correlation plots. Spearman correlation was used for correlation analysis. (d) Event free (EFS) and overall survival (OS) probability of AML patients (n=62) categorized according to the ELN2017 genetic risk classification. Rank-log test is used for statistics.

**Figure S8.**
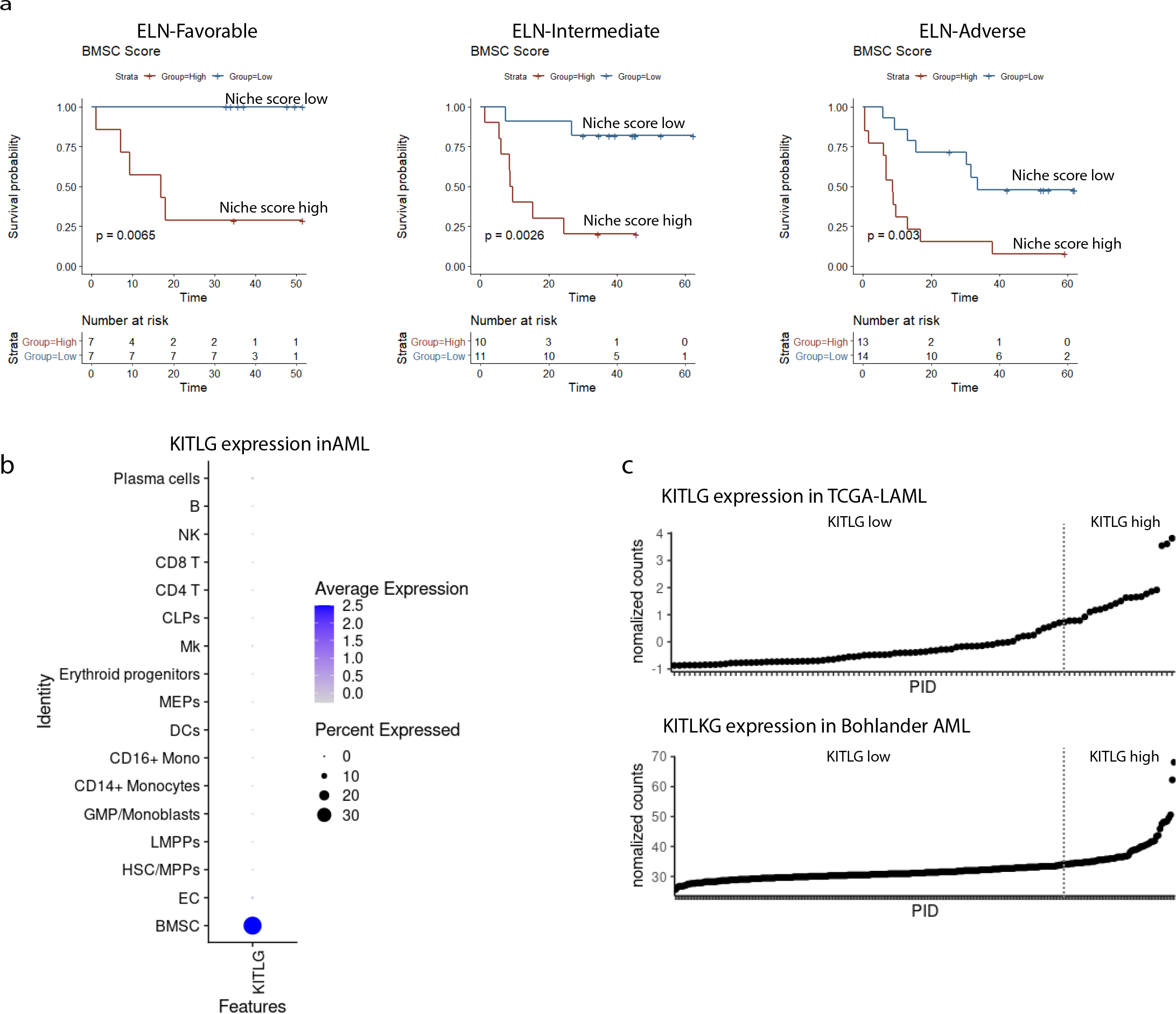
Gene signatures indicative of BMSC remodeling are associated with clinical outcome across all ELN2017 genetic risk categories. (a) Niche inflammatory remodeling and reduced KITLG expression were associated with a higher overall survival independent of ELN2017 genetic risk classification as shown by multivariate cox regression analysis in our test cohort and the TCGA-LAML validation cohort. Log-rank test is used for statistical analysis. (b) Kaplan-Meier curves for overall survival (OS) indicating improved outcome in patients with inflammatory niche remodeling (low niche score) within all ELN2017 genetic risk categories. (c) Expression of *KITLG* in bone marrow cell types in *NPM1m* AML in single cell RNAseq. Note that *KITLG* is exclusively expressed in the BMSC population. (d) *KITLG* expression in bone marrow nuclear cells as assessed in the TCGA-AML and Bohlander AML cohorts. PID, patients ID. Stratification of *KITLG* low and *KITLG* high patients was based on 75th percentile of normalized counts.

**Figure S9.**
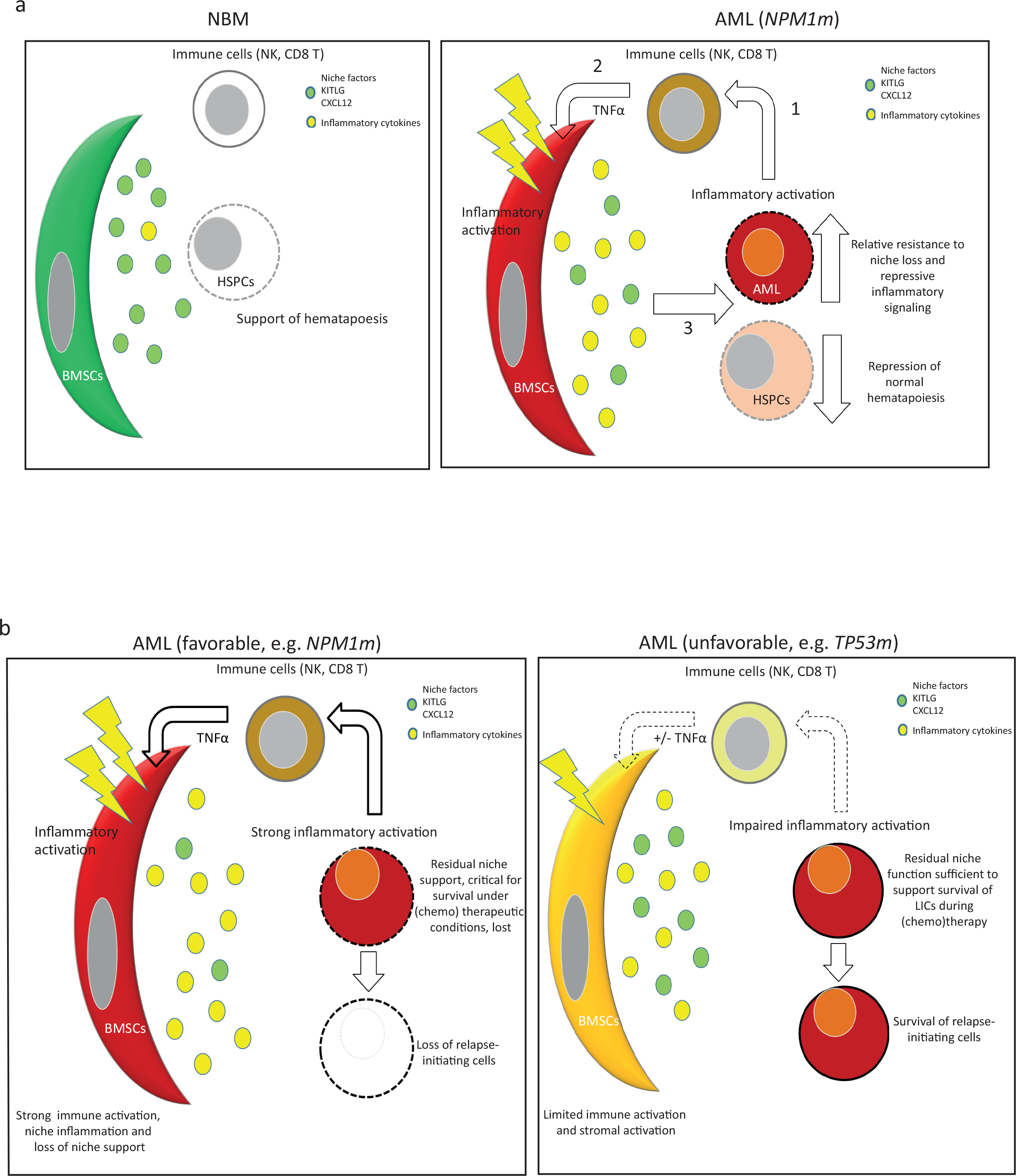
Predicted working models for the contribution of stromal remodeling to disease pathogenesis and clinical outcome in AML. a. A working model of niche inflammation mediated suppression of normal hematopoiesis and clonal advantage in AML. In AML, activation of inflammatory signaling, including immune cell activation (1) results in inflammatory activation of stromal HSPC niches (2), resulting in increased secretion of inflammatory, negative regulators of hematopoiesis and reduced expression of key HSPC maintaining factors (3). This results in the repression of normal hematopoiesis, while mutated/transformed cells display relative resistance to the repressive inflammatory signaling from HSPC niches, providing clonal advantage. b. A working model of stromal niche contributions to leukemic cell survival and outcome in AML. Leukemia (initiating) cells may remain critical dependent on residual levels of BMSC niche support (e.g. via SCF/KITLG signaling) for their survival in the context of chemotherapeutic treatment. Strong inflammation-associated loss of BMSC niches takes away this support, resulting in eradication of leukemia (relapse-initiating) cells and reduced risk of relapse. In adverse risk AML, immunogenic responses may be mitigated, ultimately resulting in relative preservation of HSPC niche support and survival of leukemia-initiating cells and increased risk of relapse in the context of chemotherapy. Note: direct effects of activated immune cells on residual leukemic cells may provide an alternative (or additional) working model for the association of niche remodeling and outcome in AML.

## SUPPLEMENTARY TABLE LEGENDS

**Supplementary Table 1. Patient characteristics**

**Supplementary Table 2. Differentially expressed genes in the HSC (low-output) cluster 0 vs. (high-output) HSC/MPP clusters 1-4**

**Supplementary Table 3. Differentially expressed genes in the NBM BMSC cluster 0 vs. other BMSC clusters**

**Supplementary Table 4. Differentially expressed genes and transcriptional programs (GSEA Hallmark) in BMSCs from AML vs. BMSCs from NBM (scRNAseq)**

**Supplementary Table 5. Differentially expressed genes in residual normal HSCs in AML vs. HSCs from NBM (scRNAseq)**

**Supplementary Table 6. Differentially expressed genes in BMSCs from AML vs. BMSCs from NBM (purified BMSC RNAseq)**

**Supplementary Table 7. Gene list reflecting the inflammatory remodeling of stromal niches in AML**

